# Dynamic noise estimation: A generalized method for modeling noise fluctuations in decision-making

**DOI:** 10.1101/2023.06.19.545524

**Authors:** Jing-Jing Li, Chengchun Shi, Lexin Li, Anne G. E. Collins

## Abstract

Computational cognitive modeling is an important tool for understanding the processes supporting human and animal decision-making. Choice data in decision-making tasks are inherently noisy, and separating noise from signal can improve the quality of computational modeling. Common approaches to model decision noise often assume constant levels of noise or exploration throughout learning (e.g., the *ϵ*-softmax policy). However, this assumption is not guaranteed to hold – for example, a subject might disengage and lapse into an inattentive phase for a series of trials in the middle of otherwise low-noise performance. Here, we introduce a new, computationally inexpensive method to dynamically infer the levels of noise in choice behavior, under a model assumption that agents can transition between two discrete latent states (e.g., fully engaged and random). Using simulations, we show that modeling noise levels dynamically instead of statically can substantially improve model fit and parameter estimation, especially in the presence of long periods of noisy behavior, such as prolonged attentional lapses. We further demonstrate the empirical benefits of dynamic noise estimation at the individual and group levels by validating it on four published datasets featuring diverse populations, tasks, and models. Based on the theoretical and empirical evaluation of the method reported in the current work, we expect that dynamic noise estimation will improve modeling in many decision-making paradigms over the static noise estimation method currently used in the modeling literature, while keeping additional model complexity and assumptions minimal.

## 1. Introduction

Computational modeling has helped cognitive scientists, psychologists, and neuroscientists to quantitatively test theories by translating them into mathematical equations that yield precise predictions [1, 2]. Cognitive modeling often requires computing how well a model fits to experimental data. Measuring this fit – for example, in the form of model evidence [3] – enables a quantitative comparison of alternative theories to explain behavior. Measuring model fit to the data as a function of model parameters helps identify the best-fitting parameters for the given data, via an optimization procedure over the fit measure (typically negative log-likelihood) in the space of possible parameter values. When fitted as a function of experimental conditions, model parameter estimation can help explain how task manipulations modify cognitive processes [5]; when fitted at the individual level, estimated model parameters can help account for individual differences in behavioral patterns [6]. Moreover, recent work has applied cognitive models in the rapidly growing field of computational psychiatry to quantify the functional components of psychiatric disorders [7]. Importantly, cognitive modeling is particularly useful for explaining choice behavior in decision-making tasks – it reveals links between subjects’ observable choices and putative latent internal variables such as objective or subjective value [8], strength of evidence [9], and history of past outcomes [10]. This link between internal latent variables and choices is made via a *policy* : the probability of making a choice among multiple options based on past and current information.

An important feature of choice behavior produced by biological agents is its inherent noise, which can be attributed to multiple sources including inattention [11, 12], stochastic exploration [39], and internal computation noise [14]. Choice randomization can be adaptive, as it encourages exploration, which is essential for learning [15]. Exploration can come close to optimal performance if implemented correctly [16, 17, 18]. However, the role of noise is often downplayed in computational cognitive models, which usually emphasize noiseless information processing over internal latent variables – for example, in reinforcement learning, how the choice values are updated with each outcome [19]. A common approach to modeling noise in choice behavior is to include simple parameterized noise into the model’s policy [2]. For example, a greedy policy, which chooses the best option deterministically, can be “softened” by a logistic or softmax function with an inverse temperature parameter, *β*, such that choices among more similar options are more stochastic than choices among more different ones. Another approach is to use an *ϵ*-greedy policy, where the noise level parameter, *ϵ*, weighs a mixture of a uniformly random policy with a greedy policy. This approach is motivated by a different intuition: that lapses in choice patterns can happen independently of the specific internal values used to make decisions. Multiple noise processes can be used jointly in a model when appropriate [20].

Failure to account for a noisy choice process in modeling could lead to under-or over-emphasis of certain data points, and thus inappropriate conclusions [21, 22]. However, commonly used policies with noisy decision processes share strong assumptions. In particular, they typically assume that the levels of noise in the policy are fixed, or “static”, with regards to some learning variable (e.g., trial for *ϵ*-greedy and value difference between choices for softmax), over the duration of the experiment, with some exceptions reviewed by [23, 24] further described in Discussion. This static assumption could hold for some sources of noise, such as computation and some exploration noise, but many other sources are not guaranteed to generate consistent levels of noise. For instance, a subject might disengage during some periods of the experiment, but not others. Therefore, existing models with static noise estimation might fail to fully capture the variance in noise levels, which can impact the quality of computational modeling.

To resolve this issue, we introduce a dynamic noise estimation method that estimates the probability of noise contamination in choice behavior trial-by-trial, allowing it to vary over time. Fig 1A illustrates examples of static and dynamic noise estimation on human choice behavioral data from [4, 5]. The probabilities of noise inferred by models with static and dynamic noise estimation are shown in conjunction with choice accuracy. In this example, choice accuracy drops steeply to a random level (0.33) around Trial 350, indicating an increased probability of noise contamination. This change is captured by dynamic noise estimation but not the static method.

**Figure 1.**
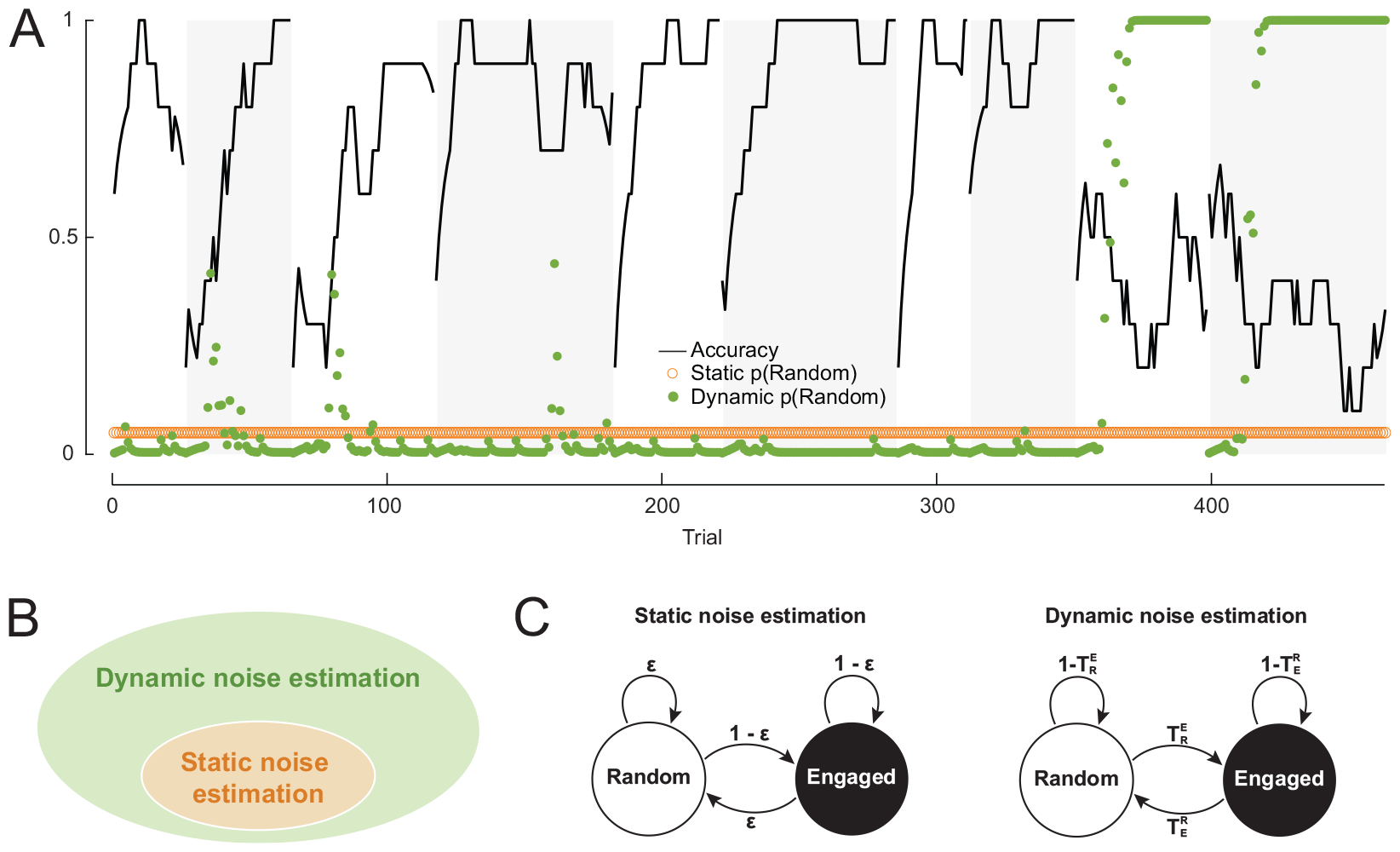
Dynamic noise estimation computes the noise levels in choices trial-by-trial. A: Example noise levels in choice behavioral data estimated by static and dynamic noise estimation methods. Background shading indicates the block design of the experiment; black line is smoothed accuracy; orange circles and green dots represent estimated static and dynamic noise levels, respectively. Data is an example subject from [4, 5]. B: Static noise estimation is a special case of dynamic noise estimation subject to an additional constraint – the static noise model space is included in the dynamic noise model space. C: Hidden Markov models representing the static and dynamic noise estimation frameworks with transition probabilities between latent states.

Our dynamic noise estimation method makes specific, but looser assumptions than static noise estimation, making it suitable to solve a broader range of problems (Fig 1B). Specifically, a policy with dynamic noise estimation models the presence of random noise as the result of switching between two latent states – the *Random* state and the *Engaged* state – that correspond to a uniformly random, noisy policy and some other decision policy assuming full task engagement (e.g., an attentive, softmax policy). We assume that a hidden Markov process governs transitions between the two latent states with two transition probability parameters, 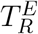 and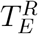, from the Random to Engaged state and vice versa. Note that static noise estimation can be formulated under the same binary latent state assumption, with the additional constraint that the transition probabilities must sum to one, making it a special case of dynamic noise estimation (see Materials and methods for proof). The hidden Markov model of dynamic noise estimation captures the observation that noise levels in decision-making tend to be temporally auto-correlated, which may be a reflection of an evolved expectation of temporally autocorrelated environments [25].

We show that noise levels can be inferred dynamically trial-by-trial in multi-trial decision-making tasks, using a simple, step-by-step algorithm (Algorithm 2). On each trial, the model infers the probability of the agent being in each latent state using observation, choice, and (if applicable) reward data. It estimates the choice probability as a weighted average of decisions generated by the Random policy and the Engaged policy, which is then used to estimate the likelihood. Therefore, dynamic noise estimation can be incorporated into any decision-making models with analytical likelihoods. Model parameters can be estimated using procedures that optimize the likelihood or its posterior distribution, including maximum likelihood estimation [26] and hierarchical Bayesian methods [27].

## 2. Modeling framework

In a multi-trial decision-making task, the agent’s data include observation-action pairs (*o*_*t*_, a_*t*_) over the learning trajectory for time *t* = 1, 2, …, *T*. In a reinforcement learning task, reward *r*_*t*_ is additionally observed on each trial. We assume that choices are generated by a Markov decision process [52]. The decision-making model leads to a policy *π*(*a*|*o*) that the agent uses to choose between discrete actions given the observation. The policy may include noise mechanisms, such as using the softmax function for action selection, and it is conditional on the model’s latent variables and parameters (e.g., learned values and learning rates for reinforcement learning models). We describe two extensions of such a decision model: the static noise estimation method that implements the classic *ϵ*-mechanism (or *ϵ*-softmax) [21] and the new dynamic noise estimation method. The parameters θ of both extended models can be optimized by maximizing the likelihood of the data given the model parameters, denoted as *ℒ* (θ). In this section, we focus only on the policy part of the models; all other model equations (such as reinforcement learning value updates) are taken from the published models and reported in Model equations.

### 2.1. Static noise estimation

Static noise policies assume that decision noise is at a constant level *ϵ* throughout the learning trajectory. At any time *t*, from the set of available actions *A*, the agent samples an action uniformly at random (with probability *ϵ*) or based on the learned policy (with probability 1 *-ϵ*). Static noise estimation can be incorporated into likelihood estimation according to Algorithm 1. Thus, any model that can be fitted with likelihood-dependent methods can incorporate static noise into its policy.

#### Algorithm 1

Static noise estimation likelihood computation

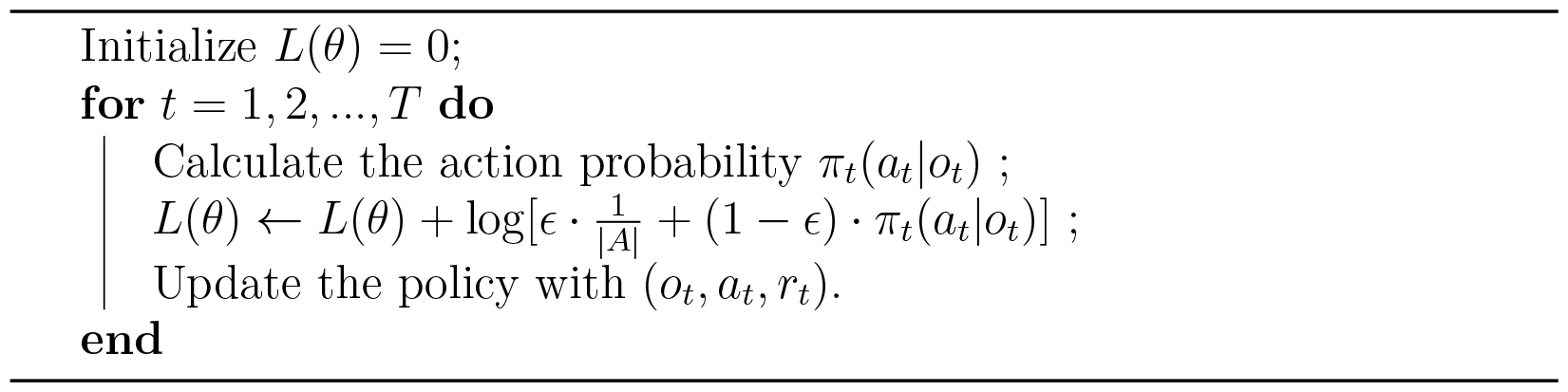

### 2.2. Dynamic noise estimation

Our dynamic noise estimation method provides a computationally lightweight procedure to estimate the trial-by-trial latent state occupancy and likelihood of the hidden Markov model described in Fig 1C. Dynamic noise estimation can be implemented according to Algorithm 2: on trial t, the likelihood, l_*t*_,

#### Algorithm 2

Dynamic noise estimation likelihood computation

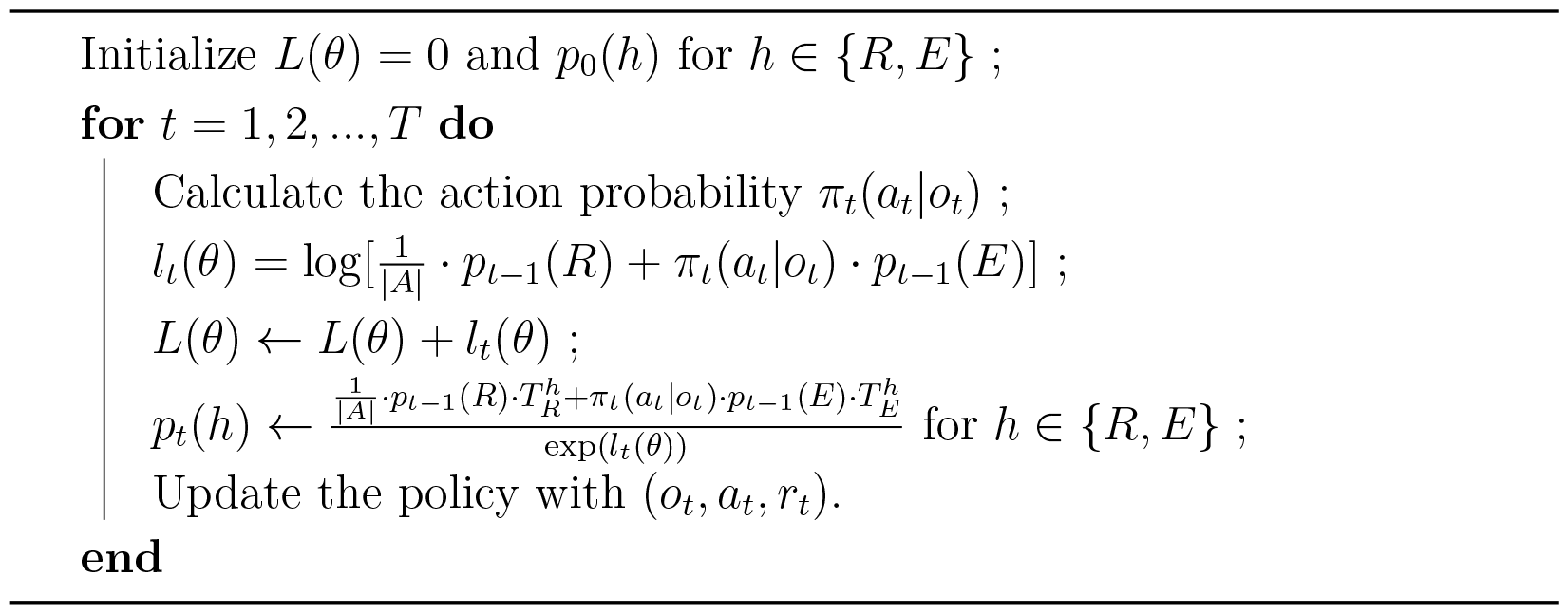

and latent state occupancy probabilities, *p*_*t*_(*Random*) and *p*_*t*_(*Engaged*), can be estimated using the observation, action, and reward data, (*o*_*t*_, a_*t*_, r_*t*_), and some engaged policy, **π**.

The full details of our dynamic noise estimation framework, which can be added on to any standard decision-making or learning model, can be found in the Materials and methods section, including the derivation of relevant mathematical equations. Here, we briefly highlight the core assumptions made by dynamic noise estimation:

1. The agent fully occupies one latent policy state on any given trial.
2. Latent state occupancy is temporally autocorrelated, and governed by a hidden Markov process: the latent state that the agent occupies on trial t conditionally depends on the latent state it occupied on trial *t* - 1.
3. Any learning involved in either latent state occurs regardless of latent state occupancy.

Additionally, the simulations and analyses below include the following non-core assumptions that can be easily modified for extended applications of our modeling framework: We assume that there are only two possible latent states, that one (“engaged”) follows the standard policy; and the other (“disengaged”) follows a uniform random policy. Both core and non-core assumptions are further discussed and explored in the discussion section.

## 3. Results

### 3.1. Theoretical benefits of dynamic noise estimation

We first performed a simulation study to demonstrate the benefits of our dynamic noise estimation approach. By definition, we expected dynamic noise estimation to explain choice data better than static noise estimation when noise levels are highly variable across trials in a temporally autocorrelated fashion. To illustrate it, we compared models implemented with static and dynamic noise estimation mechanisms on simulated data in a two-alternative, probabilistic reversal learning task widely used to assess cognitive flexibility [28], in which the correct action switched every 50 trials (Fig 2). In the simulations, we used the model with static noise to generate choice data, in which we produced periods of lapses into random behavior (e.g., due to inattention) by making the agent choose randomly between the actions.

**Figure 2.**
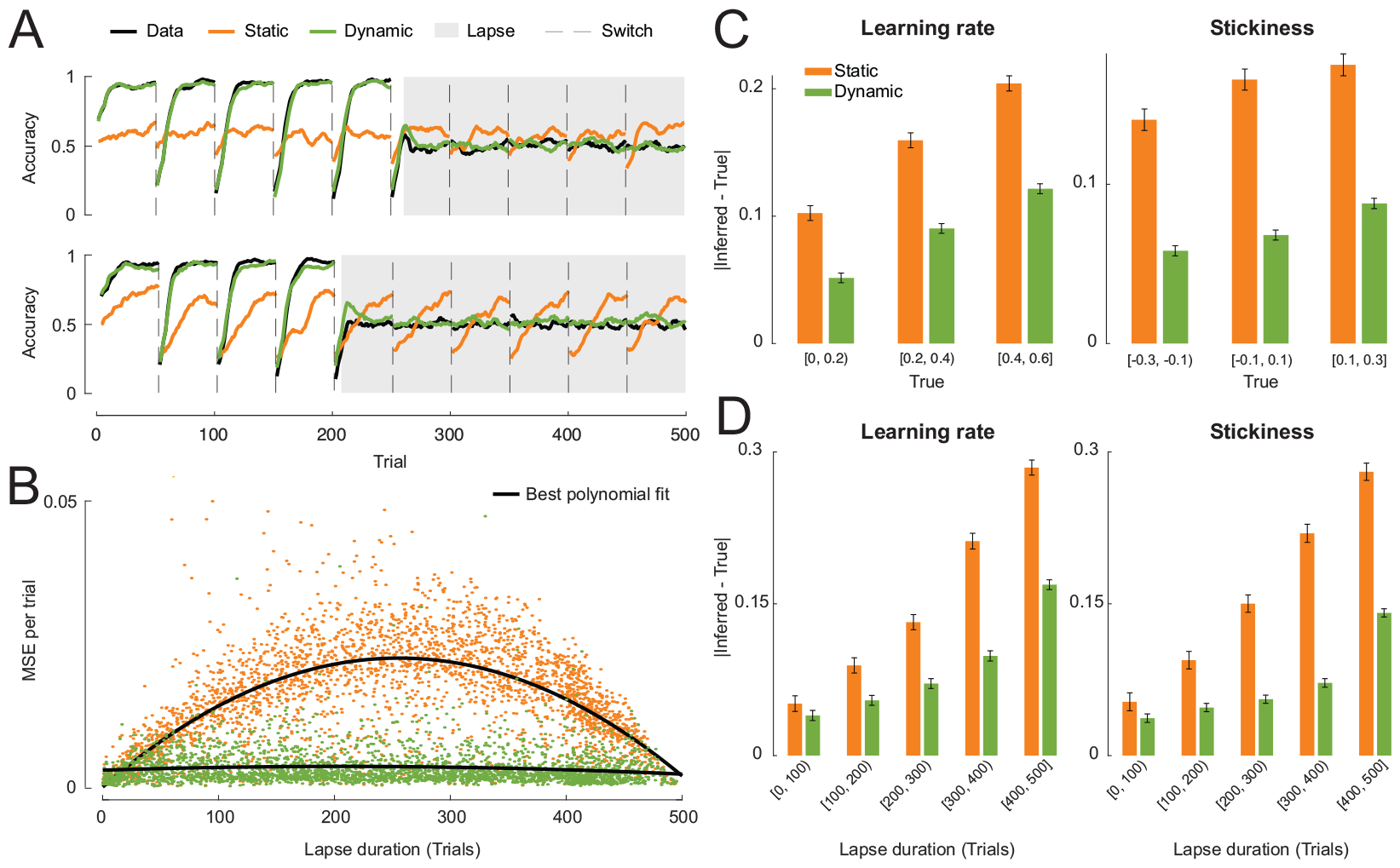
Dynamic noise estimation outperforms static noise estimation when subjects lapse into random behavior. A: Example learning curves of two simulated subjects and their best fit models with static and dynamic noise estimation; since the noise levels are fixed in the static model, the model overestimates performance in disengaged periods and underestimates it in engaged ones. B: The deviations of the best fit models’ learning curves from the data quantified by the mean squared error per trial, as a function of lapse duration. C,D: The absolute differences between the true and inferred model parameters, over true parameter value (C) and lapse duration (D).

After fitting the models to the data, we simulated behavior using the best fit parameters of both models and compared their learning curves to the data as a validation step. Fig 2A shows the learning curves of two example subjects and their best fit models. In both cases, the subjects performed at chance level (accuracy = 0.5) during lapses and better than chance otherwise. The phasic fluctuations of choice accuracy were synchronized to the reversals (dashed vertical lines). The learning curves generated by the dynamic model matched the data substantially better than the learning curves of the static model. Critically, this is true both during and outside of lapses: having to account for the lapse periods, the static noise model inferred too much noise overall, which contaminated the engaged periods. Thus, the static noise model overestimates performance in disengaged periods and underestimates it in engaged ones; by contrast, the dynamic noise model accurately captures behavior in both situations.

To further understand how the duration of lapse interacts with the effectiveness of static and dynamic noise estimation, we varied the lapse duration in the simulations. Fig 2B shows how the amounts of deviation between the learning curves of the models and data (measured by the mean squared error between the curves per trial) changed as the duration of lapse increased. Overall, the model with dynamic noise estimation was able to replicate behavior better than the static model, as the learning curves of the former matched the data more closely. Although lapses only weakly affected the fit of the dynamic noise model, the static model fitted worse in the presence of lapses, especially when lapse and non-lapse periods were intermixed in the learning trajectory.

Next, we tested how well the true parameters used to generate the data could be recovered by the static and dynamic models (Fig 2C). Both learning parameters (learning rate and choice stickiness) were better recovered by the dynamic model, as measured by the absolute amounts of differences between the true and recovered (best fit) parameters. The advantage of the dynamic model in parameter recovery persisted over the whole range of parameter values sampled in the simulations and various lengths of lapses, with weaker effects when lapses were short relative to the duration of the experiment (less than 20%). Additionally, we performed the same set of analyses using the static model as the ground truth (Fig A.7). As expected, overall, the static model outperforms the dynamic model, even though both models can accurately capture behavior and recover true parameter values, since the dynamic model space fully includes the static models.

To verify that including dynamic noise estimation would not undermine a model’s robustness, we performed validation and recovery analyses on data simulated with the dynamic noise model in the same probabilistic reversal task environment used in the previous simulations. In model validation, the dynamic model reproduced behavior more closely than the static model in both the engaged state and the random state: the dynamic noise model showed much more sensitivity to the latent state than the static noise model. (Fig 3A). This suggests that fitting a model with static noise estimation when the underlying noise mechanism of the data is dynamic could lead to inaccurate interpretations of the behavior and model.

**Figure 3.**
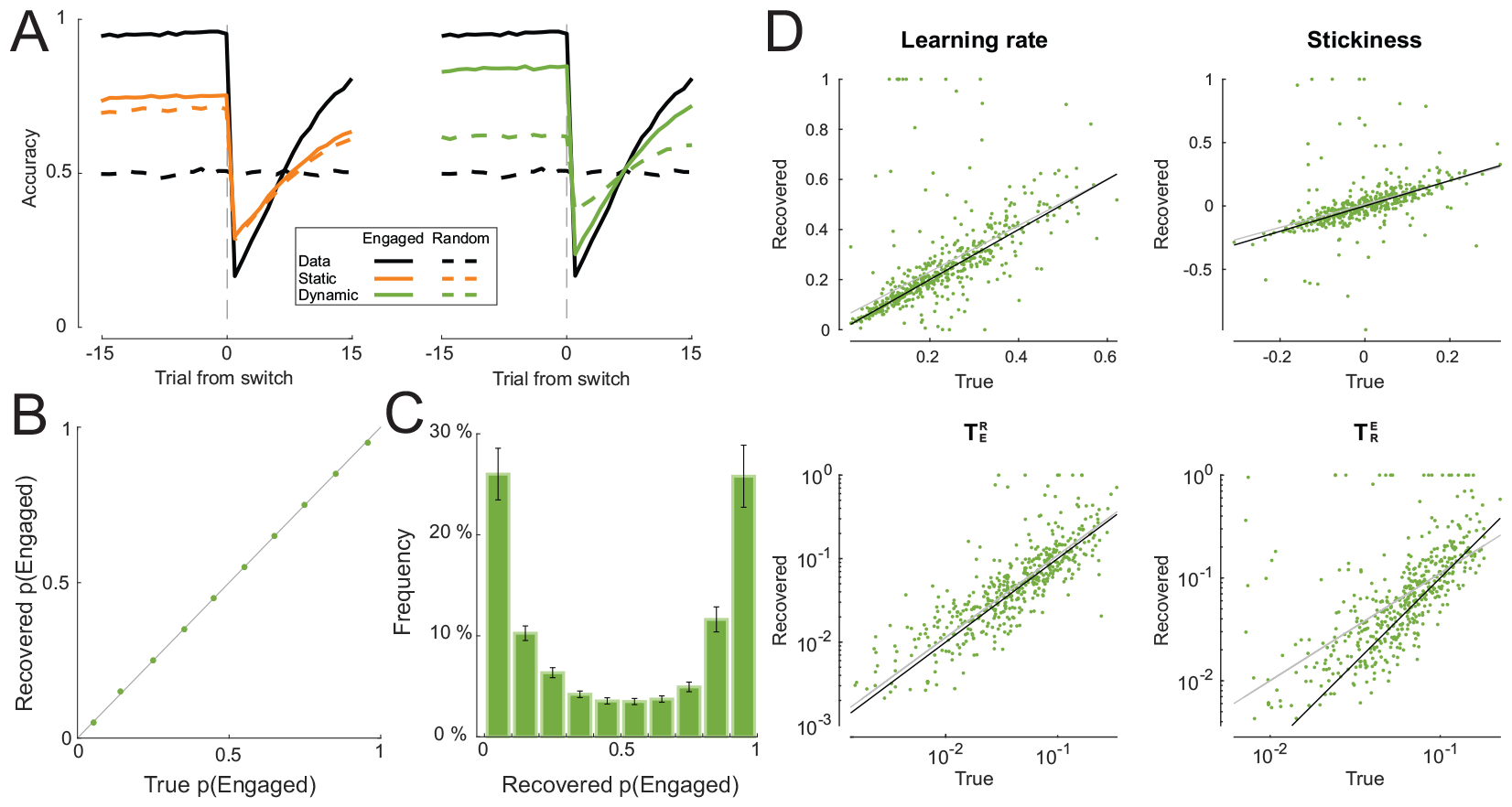
The dynamic noise estimation model validates and recovers robustly. A: Validation of best fit models with static and dynamic noise estimation against simulated data using learning curves around switches for both Engaged and Random trials. B: The recovered occupancy probability of the Engaged state, *p*(*Engaged*), over the true occupancy probability used to simulate the data. C: The distribution of the recovered occupancy probability. D: Recovered model parameters against their true values. In each plot, the black line is the least squares fit of the points and the grey line is the identity line for reference.

Furthermore, we confirmed that the occupancy probabilities of the latent states and model parameters were recoverable by fitting the dynamic model to the simulated data to infer the quantities of interest. The occupancy probability of the Engaged state, *p*(*Engaged*), was perfectly recovered across its range of values (Fig 3B). The inferred or recovered values of *p*(*Engaged*) formed a symmetric, bimodal distribution with peaks near 0 and 1, suggesting that both latent states were visited equally frequently and that the model was confident, for the majority of the time, that the agent was in either latent state (Fig 3C). The true values of all model parameters were recoverable through fitting (Fig 3D).

### 3.2. Empirical evaluation of dynamic noise estimation

The above analyses based on controlled simulations showed that, theoretically, dynamic noise estimation could substantially improve model fit and parameter estimation, especially in the presence of prolonged lapses. We next tested the method on empirical datasets to verify whether and to what extent this conclusion stands when the data is collected from real animal and human subjects while the true generative model is unknown. To help set fair expectations for the applications of dynamic noise estimation in practice, we thoroughly evaluated the method on four published datasets featuring diverse species, age groups, task designs, behaviors, cognitive processes, and computational models. Table 1 summarizes the population, task, and model information about these datasets.

**Table 1:** Summary of empirical datasets.

| Dataset | Population | Task | Model |
| --- | --- | --- | --- |
| Dynamic Foraging [29] | Mice | Two-armed bandits with probabilistic reversal | Reinforcement learning with dynamic learning rates |
| IGT [30] | Young and old adult humans | Iowa gambling task | A hybrid of exploitation and exploration processes [31] |
| RLWM [32] | Adult humans | Reinforcement learning and working memory | A hybrid of reinforcement learning and working memory processes |
| 2-step [33] | Developing and adult humans | Two-step task | A hybrid of model-based and model-free learning processes |

For each dataset, we used either the winning model in the original research article or an improved model from later work. We implemented and compared two versions of each model: one with static noise estimation and one with dynamic noise estimation. The models were fitted on each individual’s choice data using maximum likelihood estimation for simplicity, although the noise estimation methods are both also compatible with more complex likelihood-based fitting procedures. The fitted models were compared using the Akaike Information Criterion (AIC) [34], since it yielded better model identification than the Bayesian Information Criterion (BIC; Fig A.8). Fig 4 shows the model-fitting results at both the individual and group levels, as well as the absolute percentage of fit improvement, using the fit measure of negative log-likelihood (NLLH), made by applying dynamic noise estimation instead of static noise: 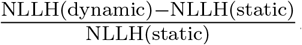.To compare the models at the group level, we report the p-values of one-tailed Wilcoxon signed-rank tests with the alternative hypothesis that the AIC values of the dynamic model were lower than those of the static model. Additionally, we report the protected exceedance probability (pxp) [35] of the dynamic model. At the group level, dynamic noise estimation significantly improved model fit compared to static noise estimation on the Dynamic Foraging (ΔAIC = *-*8.31, p = 0.0002, pxp = 0.96) and IGT (ΔAIC = *-*2.79, p = 3.48 *×* 10^*-*12^, pxp = 1.00) datasets. This populational difference was present but not statistically significant on the RLWM (ΔAIC = *-*1.43, p = 0.83, pxp = 0.38) and 2-step (ΔAIC = *-*3.04, p = 0.47, pxp = 0.44) datasets. While the absolute percentage of fit improvement is small for most subjects, it can be very high for some, which may enable researchers to still include “noisy” subjects in their analyses without biasing results (median = 0.29% for Dynamic Foraging, 1.21% for IGT, 0.16% for RLWM, and 0.3% for 2-step). Since static noise estimation is fully nested in dynamic noise estimation, the absolute fit improvement by dynamic noise estimation is strictly positive.

**Figure 4.**
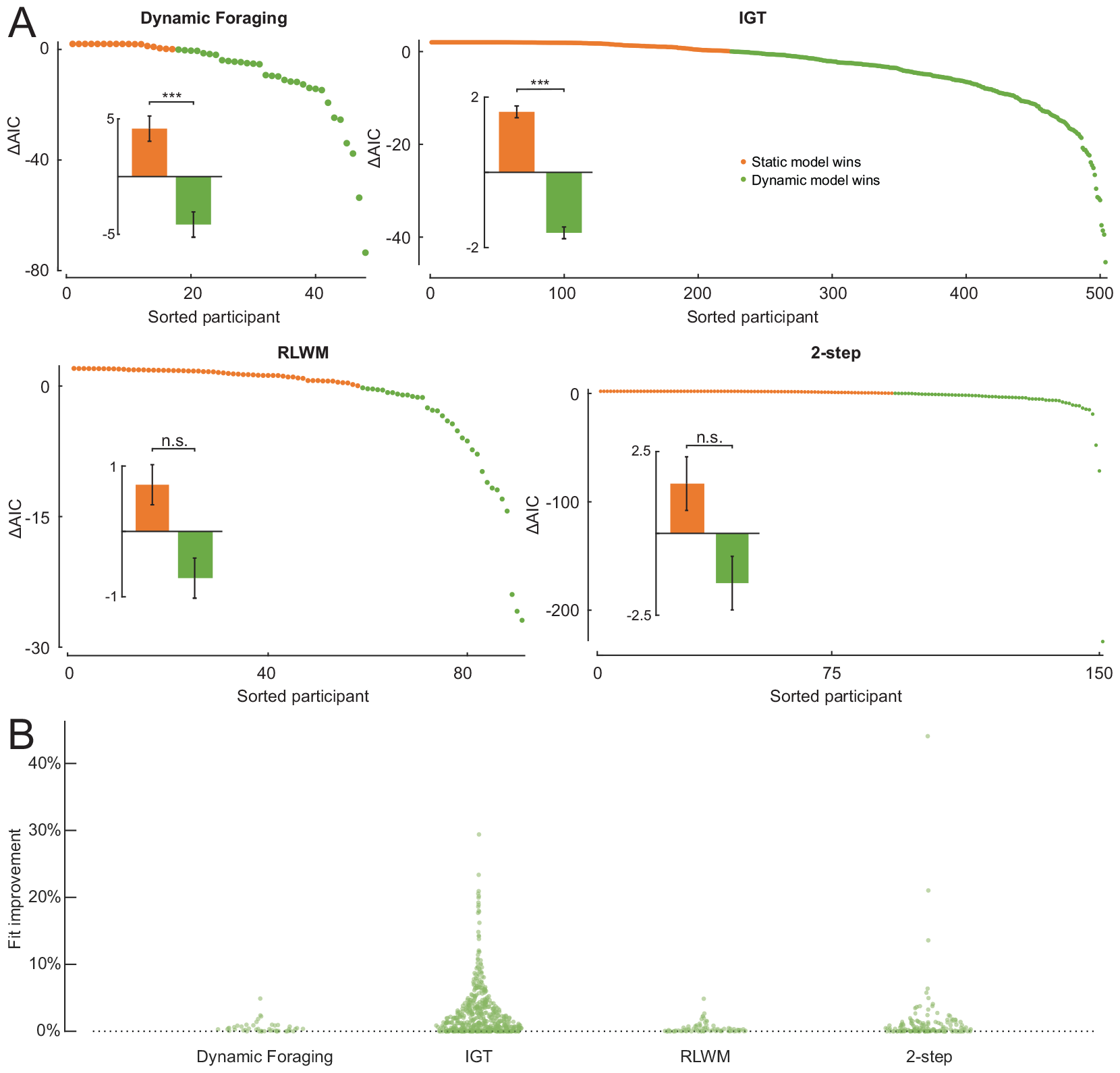
Dynamic noise estimation can improve model fit on empirical data. A: Evaluation of model fit on four empirical datasets based on the AIC. In each panel, the plot shows the difference in AIC for each individual between the models with static and dynamic noise estimation mechanisms. A positive value (orange) indicates that the static model is favored and a negative value (green) means that the dynamic model is preferred by the criterion. The inset shows the mean difference in AIC between the models at the group level. Significance levels are defined as *\*\* \** if p*<* 0.001, *\*\** if p*<* 0.01, *\** if p*<* 0.05, and n.s. otherwise. B: The absolute percentage of improvement on fit, measured by the negative log-likelihood, by dynamic noise estimation from static noise estimation.

As detailed in Materials and methods, the likelihood of the dynamic noise estimation model should not be worse than that of the static model, since the latter is equivalent to a special case of the former. This relationship was confirmed by the fitting results on all four empirical datasets: for individuals whose data were better explained by the static model, the ΔAIC values were upper-bounded by 2, which corresponded to the penalty incurred by the extra parameter in the dynamic model. In other words, the dynamic model did not impair likelihood estimation in practice, which aligned with our prediction.

We additionally validated both models against behavior and found no significant differences between the static and dynamic noise models (Fig A.9). We verified that the quantities specific to dynamic noise estimation, including the occupancy probability and noise parameters, were recoverable (Fig A.10). The distributions of the estimated occupancy probability of the Engaged state, *p*(*Engaged*), were heavily right-skewed and long-tailed. This indicates a scarcity of data in the Random state overall, which likely led to a lack of transitions from the random state to the engaged state and, thus, under-powered the recovery of 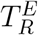, causing it to be noisier than the recovery of 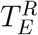.

Knowing that likelihood favors the dynamic model over the static model, the remaining questions are: *how* does this improvement manifest, and does it impact the insights we can gain from computational modeling? To address these questions, we compared the values of best fit parameters between both models (Fig 5). On the Dynamic Foraging dataset, the values of the positive learning rate and forgetting rate parameters, which govern the value updating rate of rewarded actions and the forgetting rate of unchosen actions (see Model equations for the full model description), increased at the group level (two-tailed Wilcoxon signed-rank test p = 7.56 *×* 10^*-*7^ for positive learning rate and p = 2.66 *×* 10^*-*5^ for forgetting rate). We speculate this may suggest that dynamic noise estimation helped the model capture faster learning dynamics in the task, which may have led to the improved fit. On the RLWM dataset, the distributions of the bias (p = 0.0016) and stickiness (p = 0.0022) parameters, which represent the bias in learning rate for unrewarded actions compared to rewarded actions and the choice stickiness (see Model equations for the full model description), both shifted in the positive direction. On the 2-step dataset, the softmax inverse temperature parameter for the second-stage choice was also estimated to increase after incorporating dynamic noise estimation into the model (p = 8.8 *×* 10^*-*6^). Similarly, on the IGT dataset, the softmax inverse temperature parameter increased significantly (p = 2.78 *×* 10^*-*7^). An increase in the inverse temperature parameter can be interpreted as capturing a policy that is less noisy and more sensitive to internal variables; these results highlight the success of the dynamic noise model in identifying noisy time periods and decontaminating on-task periods from their influence.

**Figure 5.**
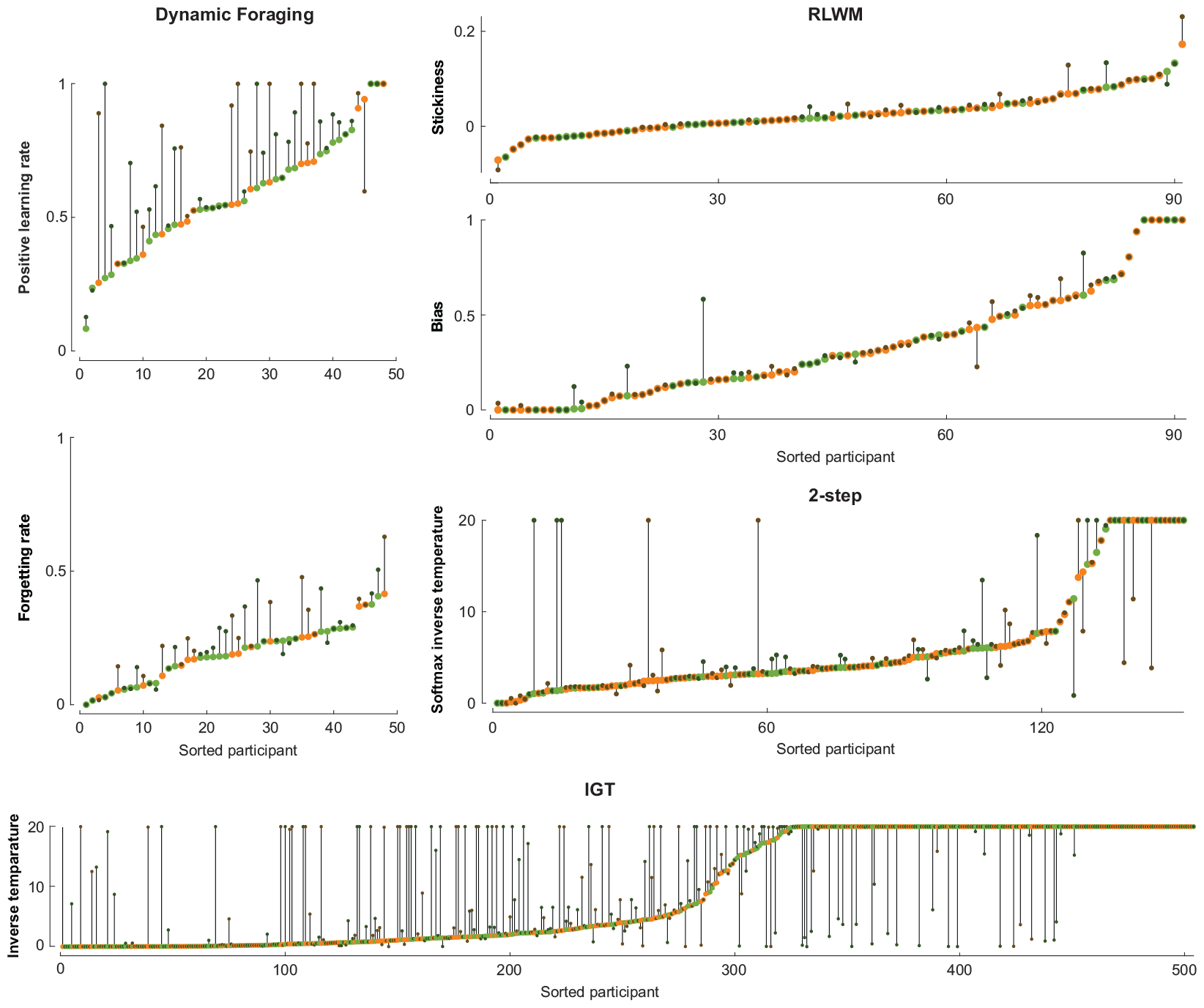
Dynamic noise estimation can lead to shifted parameter fit. Changes in best fit parameter values between the models with static and dynamic noise estimation mechanisms for each individual. Individual data points are color-coded according to the winning model by AIC: orange if the static model fitted better and green if the dynamic model fitted better.

Besides the policy parameters, the noise parameters also showed distributional differences that were correlated with improved fit. Fig 6 illustrates the relationship between the static noise parameter, *ϵ*, and the dynamic noise parameter, 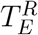, on all four empirical datasets. For individuals whose data were better explained by the static noise model according to the AIC, 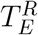 and *ϵ* were estimated to take on comparable and highly correlated values (Dynamic Foraging: Kendall’s τ = 0.84, p = 5.67 *×* 10^*-*5^; IGT: τ = 0.82, p = 1.23 *×* 10^*-*67^; RLWM: τ = 0.89, p = 6.78 *×* 10^*-*23^; 2-step: τ = 0.84, p = 1.42 *×* 10^*-*26^). This observation was in line with our expectation: when the static model was favored by the AIC, the difference in likelihoods between both models must be smaller than the penalty incurred by the extra parameter in the dynamic model (2 for AIC), which means both models fitted similarly to the data. On the other hand, when the dynamic model outperformed the static model, 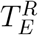 was estimated to be lower than *ϵ* (Dynamic Foraging: one-tailed Wilcoxon signed-rank test p = 0.031; IGT: p = 4.90 *×* 10^*-*8^; RLWM: p = 0.0072; 2-step: p = 0.0017). A similar, though noisier, relationship between 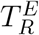 and 1 *-ϵ* was also observed on all empirical datasets (Fig A.11). No consistent strong correlations were found across datasets between the noise parameters of the dynamic model (softmax inverse temperature, 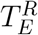, and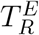; Fig A.12). The lower values of the dynamic noise parameter than the static noise level parameter, which is the average noise level, indicate that the dynamic model successfully separated noisy trials from engaged trials.

**Figure 6.**
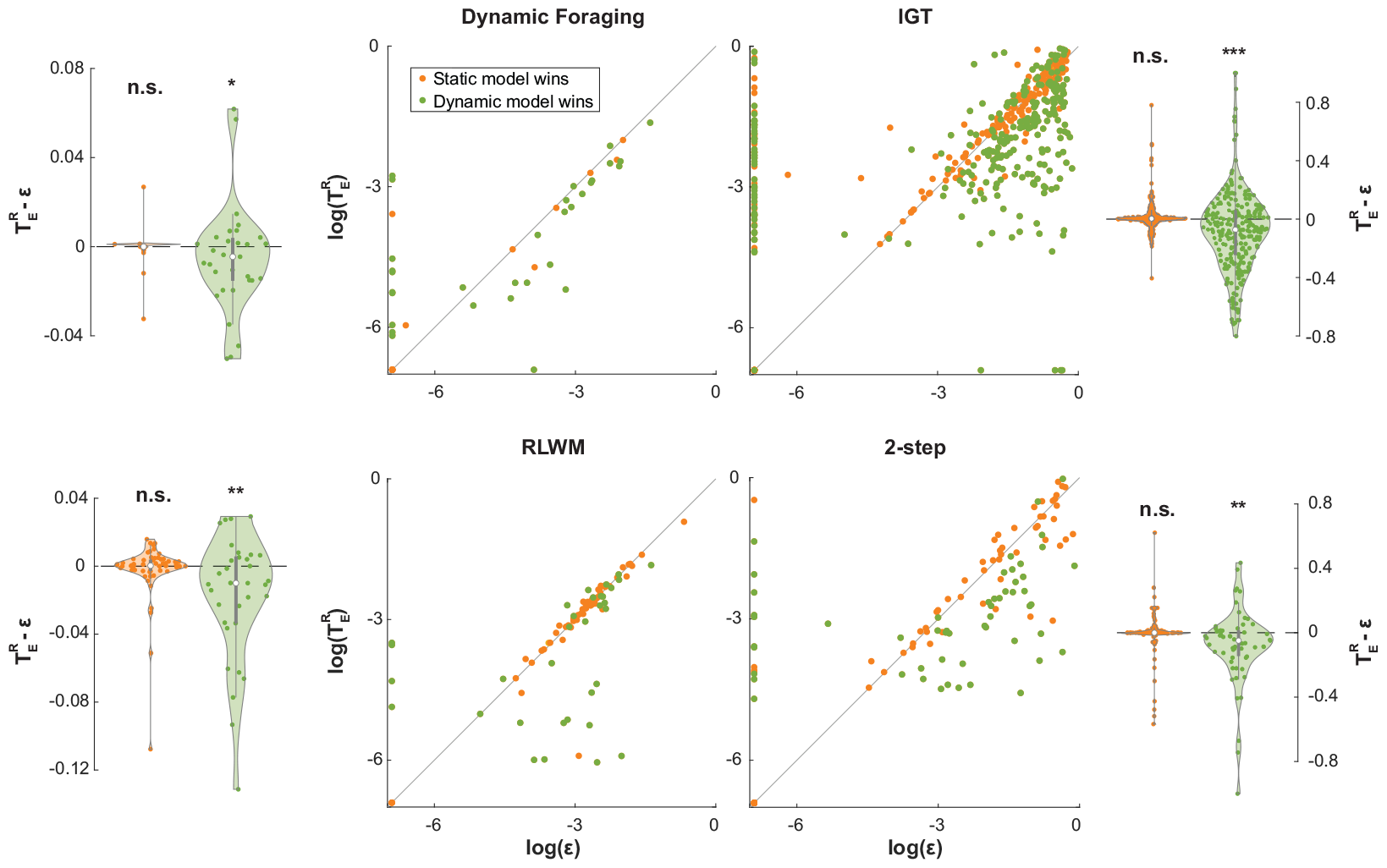
Improved fit by dynamic noise estimation is correlated to decreased noise parameter estimates. The dot plots in the center illustrate the relationship between the best fit dynamic and static noise parameters (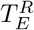and *ϵ*) on log scale, with each dot representing an individual. The violin plots on the sides show the differences between the best fit dynamic noise parameter, 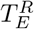, and static noise parameter, *ϵ*, at the individual and group levels.

To demonstrate the behavioral relevance of the latent state occupancy predicted by dynamic noise estimation, we investigated whether behavior differed between the putatively engaged and lapsed trials (as identified by our approach) on four empirical datasets: Dynamic Foraging [29], IGT [30], 2-step [33], and RLWM [4, 5] (Fig A.13). In general, we found that behavior shifted towards random patterns from engaged trials to lapsed trials. Interestingly, some components of behavior regressed to randomness more than others. For example, on the IGT dataset, behavioral changes were driven by decks A and D, but not decks B and C. On the RLWM dataset, the win-stay probability decreased more than the lose-shift probability across set sizes. Lapses identified by dynamic noise estimation varied in lengths and occurred throughout learning, with no strong evidence for consistently more frequent lapses in specific parts of the experiments across datasets (Fig A.14).

Furthermore, we related the estimated latent state occupancy to an independent measure of behavior – reaction time – using regression analyses on both the group and individual levels on two empirical datasets with published reaction time data: RLWM [32] and 2-step [36]. On both datasets, we found significant inverted-U relationships between reaction time and *p*(*Engaged*) both between- and within-individual (Fig A.15). The squared average reaction time inversely predicted the average *p*(*Engaged*) across participants 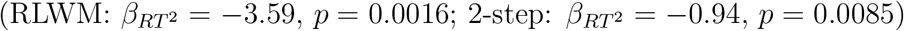. We found a similar relationship within-participant across trials while accounting for a random effect of participant identity 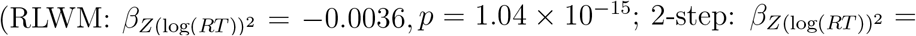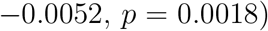. These results suggest that low task engagement estimated by dynamic noise estimation is more likely to occur in trials with unusually short and long reaction time, which potentially includes when participants answer excessively fast due to boredom or very slowly due to external distraction, such as multitasking.

## 4. Discussion

Our results show that dynamic noise estimation can improve model fit and parameter estimation both theoretically and empirically, qualifying it as a candidate alternative to static noise estimation, despite one additional model parameter. Our approach is especially powerful and effective in the presence of lapses, since it explains more variance in the noise levels of choice behavior. Additionally, it is generalizable and versatile: it can be applied to any decision policies with analytical likelihoods and be incorporated into any likelihood-based parameter estimation procedures, making it an accessible and computationally lightweight extension to many decision-making models. Another benefit of dynamic noise estimation is that it could help avoid excluding whole individuals or sessions due to poor performance, thus improving data efficiency. Dynamic noise estimation takes effect by identifying periods of choice behavior that are better explained by random noise than the learned policy (e.g., lapses). The likelihoods of these noisy periods are lower-bounded by that of the random policy, which limits the impacts of these trials on the estimation of the overall likelihood and model parameters. Thus, dynamic noise estimation can mitigate the effects of noise contamination on model-fitting. On the contrary, static noise estimation does not provide a meaningful lower bound to the likelihood of noisy data, such that relatively noisy parts of the behavior may heavily bias parameter estimation.

Thus, using dynamic instead of static noise estimation could allow fewer individuals to be excluded due to noisy behavior. For example, without dynamic noise estimation, the last two blocks in Fig 1A might lead to the exclusion of this subject by some performance-based criterion. However, dynamic noise estimation might allow fitting of the whole individual’s data with minimal contamination due to the noisy blocks, even though it may not improve modeling dramatically for most participants. This outcome can be particularly desirable when data collection is challenging or expensive, such as in clinical populations, neuroimaging experiments, and time-consuming tasks.

Although the putative lapses identified by dynamic noise estimation may correlate with lower choice accuracy, dynamic noise estimation has a number of advantages over approaches that rely solely on accuracy to identify lapses. First, when more than one action is available, dynamic noise estimation can use information in both the correctness and the choice identities to estimate lapse rates. As a result, it can distinguish random behavior from non-random components of decision-making such as learning and bias, which might drive the accuracy to the random level. Second, dynamic noise estimation accounts for the temporal autocorrelation of noise between trials, which is characteristic of lapses, by factoring noise information from previous trials in predicting the noise level of the next trial. Indeed, Fig A.16 shows that the probability of lapsing is not directly related to degree of accuracy. Third, the application of dynamic noise estimation is independent of the task design: it does not require task-specific tuning of any hyper-parameters or criteria.

Other approaches have been proposed to consider non-static noise or exploration, including models where noise parameters evolve trial-by-trial. For example, some decision models with softmax policies allow decision certainty to increase over learning, by defining the inverse temperature parameter or the value difference between choices as a parameterized function of time or certainty [37, 38, 39]. While these models may help capture the decrease in choice randomization over the experiment, they can only account for decision noise that changes in an incremental fashion (e.g., gradually decreasing), but not lapses that could occur unexpectedly throughout the experiment. Our approach instead relies on the assumption that participants may switch between finite, discrete late states abruptly, which is supported by behavioral findings for discrete policies [40, 41].

Biologically, our latent state assumption aligns with an established literature on how norepinephrine modulates attention, a major contributor to varying noise levels: the phasic or tonic mode of activity of the noradrenergic locus coeruleus system closely correlates to good or poor task performance [42, 43]. It is worth noting that the binary assumption of the latent states may not always be accurate. Nonetheless, it is a less strict assumption than that of static noise estimation, which additionally assumes that the probability of transitioning into each latent state is independent of the current state. Thus, although dynamic noise estimation may be limited by its binary latent state assumption, it is still more suitable to solve a broader range of problems than static noise estimation.

Compared to other recent work identifying discrete latent policy states, namely the GLM-HMM model [44], dynamic noise estimation has the advantages of simplicity, accessibility, and versatility. Contrary to our method, GLM-HMM additionally assumes that all decision policies can be described as generalized linear models, which limits its applications to descriptive models rather than cognitive process models. The parameter estimation procedure for GLM-HMM does not generalize trivially when this assumption is challenged (e.g., with process models such as reinforcement learning). On the other hand, our likelihood estimation procedure for dynamic noise estimation can be readily plugged into any existing likelihood-based optimization procedure to fit both descriptive models and process models.

We recommend that the user keep in mind the assumptions outlined in the beginning of the Results section when applying our modeling framework to their data. Dynamic noise estimation can be applied to any multi-trial decision-making tasks and models with analytical likelihoods, especially when more than one action is available in the task. Assumption 3 (the latent state only affects the policy, but not the underlying process) imposes a limitation to our approach: in the random state, information is still being processed (e.g., action value updating), but not used for decision-making. Removing this assumption can significantly complicate the inference process over the latent state by making the likelihood intractable, and thus making the inference process much less accessible. Addressing this limitation will be an important direction for future work.

Other non-core assumptions of the method may appear as limitations, but can be easily extended, such as the nature of the engaged and disengaged policies and even the number of states itself. For example, an extension to the likelihood estimation procedure derived in the current work is to apply it on policy mixtures in a broader sense – i.e., hidden Markov models that involve two or more latent states of any eligible policies – rather than a fixed random policy and some other decision policy (e.g., softmax) as presented in the current work. This extension allows us to fit mixture models between two or more decision policies to capture the switching between different strategies. When applying our framework to fit such mixture models, we recommend that the user check Assumption 1 (the agent fully occupies a single latent decision state), as it may not be appropriate for all mixture models. For example, the RLWM model [4] is a mixture of a reinforcement learning process and a working memory process, which could technically be modeled as two latent policy states. However, Assumption 3 is biologically implausible here: participants are unlikely to transition from fully occupying one policy state to the other between trials since reinforcement learning and working memory operate concurrently.

Future work should also further validate dynamic noise estimation experimentally, for example, by comparing estimated occupancy probabilities to an independent measure of attention or task-engagement and testing whether inferred latent states capture this measure. Possible approaches include to measure task-engagement based on choice behavior [45], reaction time [46], pupil size [47], and event-related brain potentials [48]. If the occupancy probability can indeed serve as an objective measure of attention to the task, it could be applied to behaviorally characterize attentional mechanisms in computational psychiatry [49], especially for patients with attention-deficit/hyperactivity disorder (ADHD) [50]. Another potential future direction is to explore whether dynamic noise estimation changes the interpretations of behaviors and models when applied to other decision policies than the softmax policy, such as Thompson sampling [17] and the upper confidence bound algorithm [51].

In conclusion, our dynamic noise estimation method promises potential improvements over the static noise estimation method currently used in the modeling literature of decision-making behavior. Dynamic noise estimation enables us to capture different degrees of task-engagement in different task periods, limiting contamination of model-fitting by noisy periods, without requiring ad-hoc data curating. Based on the theoretical and empirical evaluation of the method reported in the current work, we expect that dynamic noise estimation in modeling choice behavior will strengthen modeling in many decision-making paradigms, while keeping additional model complexity and assumptions minimal.

## 5. Materials and methods

### 5.1. Mathematical formulation of dynamic noise estimation

The dynamic noise estimation method models decision noise by assuming that the agent is in one of two latent states at any given time: the *random state* in which the agent chooses actions uniformly at random or the *engaged state* in which decisions are made according to the true model policy. The transitions between both states are governed by two parameters: 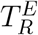 and 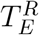, the probabilities of transitioning from the random state to the engaged state and vice versa. From these transition probabilities, we can calculate the stay probability for each latent state: 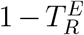 for the random state and 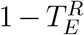for the engaged state.

The state is composed of an observation *o*_*t*_, often encoding the stimulus, and unobserved, latent variables including the learned policy and *h*_*t*_, where *h*_*t*_ ∈ {*R, E*} indicates whether the agent is in the random state or engaged state at time *t*. It is further assumed that *r*_*t*_ and *o*_*t*_ are conditionally independent of the latent states up to time t given the observed data history, since rewards and future observations in behavioral experiments do not depend on subjects’ unobserved mental states.

Our goal is to maximize the following log-likelihood:

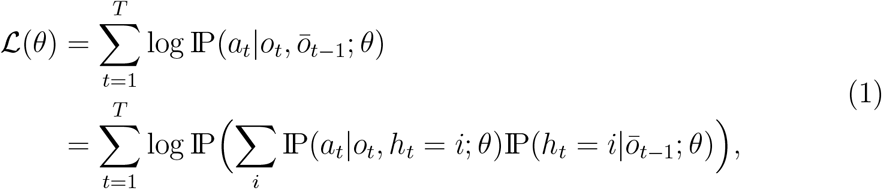

where *ō*_*t-*1_ denotes the observation-action-reward triplets up to time *t* - 1. The probability on the right of Eq 1, the occupancy probability of the latent state *i* ∈ {*R, E*} at time *t*, is not trivial to compute. Denoting it as *p*_*t*_(*i*), we have

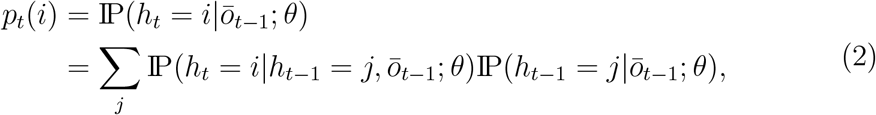

where *j* ∈ {*R, E*} and

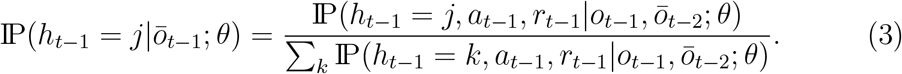

Notice that for any given *k*, each term in the denominator of the right-hand side of Eq 3, as well as the nominator with *k* = *j*, is equal to

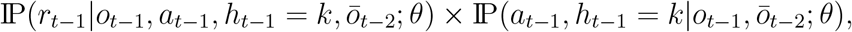

the first term of which is independent of *h*_*t-*1_ and is, therefore, canceled out between the nominator and denominator in Eq 3. Thus,

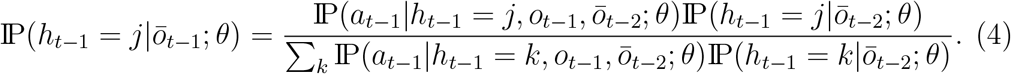

We can now compute *p*_*t*_(i) by plugging Eq 4 into Eq 2, which then allows us to calculate *ℒ* (θ) by plugging Eq 2 into Eq 1. The probabilities needed to infer *p*_*t*_(i) and *ℒ* (θ) can be iteratively updated according to Algorithm 2 over the learning trajectory. These calculations can be easily incorporated into fitting procedures based on optimizing the model’s likelihood, including maximum likelihood estimation and hierarchical Bayesian modeling.

#### 5.1.1. The relationship between static and dynamic noise estimation

Static noise estimation can be formulated under the binary latent state assumption of dynamic noise estimation (Fig 1B), with the additional constraint that the probability of transitioning into each latent state is independent from the current state:

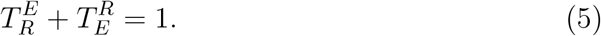

In other words, the probabilities of transitioning to the random state from the engaged state must be equal to the probability of transitioning to the random state from the random state:

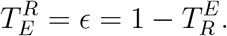

Similarly, the probabilities of transitioning into the engaged state from the random state and the engaged state must be equal:

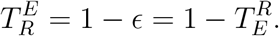

Both the above relationships can be summarized by Eq 5.

Therefore, static noise estimation is a special case of dynamic noise estimation with an additional assumption described by Eq 5, as illustrated in Fig 1C. It can also be experimentally verified that dynamic noise estimation converges to static noise estimation once this constraint is added to the model-fitting procedure (results not included).

Theoretically, with optimal parameters, the likelihood estimates made by the dynamic noise estimation model must be no worse than those made by the static noise estimation model. In practice, this relationship may not hold if the optimizer fails to converge to the global minimum when fitting the dynamic model. However, this issue can be circumvented by initializing the parameter values of the dynamic model to the best fit parameters of the static model (e.g., 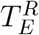 as 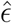 and 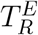 as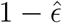).

#### 5.1.2. Initializing p(Engaged)

In the above formulation, the starting points of the estimated latent state occupancy probabilities, *p*(*Engage*d) and *p*(*Random*) = 1 *-p*(*Engaged*), are undefined, since dynamic noise estimation is compatible with any valid (staying engaged), and initial values of these probabilities. Therefore, the user can choose the most appropriate initial *p*(*Engaged*) for their data. Some potential candidates, reflecting different assumptions, include: 1 (initially engaged), 0.5 (equal chance of either), 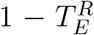 (staying engaged), and 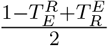(average noise level). Alternatively, the initial *p*(*Engaged*) value can be fitted as a free parameter, which may reduce bias in the estimation of latent state occupancy, but at the cost of increased model complexity. All models in the current work are fitted with initial 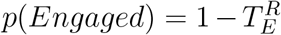, which ensures that the dynamic noise model fully includes the static model, since *p*(*Engaged*) of the static model is always 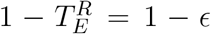. For reference, in Figure A.16, we show the estimated *p*(*Engaged*) trajectories for different initialization methods on the RLWM dataset. This indicates that differences in initialization lead to differences only in the very first few trials of a learning block.

### 5.2. Analysis methods

#### 5.2.1. Simulation setup

The task environment in which the data were simulated for the theoretical analyses had two alternative choices with asymmetrical reward probabilities (80% and 20%) that reversed every episode. Each agent was simulated for 10 episodes with 50 trials per episode. The simulations with lapses included data from 3,000 individuals generated by the model with the static noise mechanism (Fig 2). Model parameters were sampled uniformly between reasonable bounds: learning rate *∼* Uniform(0, 0.6), stickiness *∼* Uniform(*-*0.3, 0.3), and *ϵ ∼* Uniform(0, 0.2). For each individual, we simulated a lapse into random choice behavior whose duration was sampled uniformly at random between 0 and the length of the experiment (500 trials). During the lapse, the agent was forced to randomly choose between the two available actions. In the analyses shown in Fig 3, we simulated data of 1,000 individuals using the model with the dynamic noise mechanism. The parameters were sampled from the following distributions: learning rate *∼* Beta(3, 10), stickiness *∼* Normal(0, 0.1), 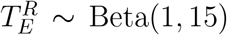, and 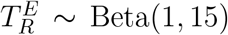*∼* Both models were fitted to the simulated data per individual.

#### 5.2.2. Empirical datasets and models

All empirical data were downloaded from sources made publicly available by the authors of the corresponding research articles. The data of all individuals were included except that for the IGT dataset [30], we selected for the studies that used the 100-trial versions of the task. For the Dynamic Foraging (n=48) [29] and 2-step (n=151) [33] datasets, the winning models from the original papers were used in our analyses. Since the article containing the IGT dataset (n=504) [30] did not report modeling results, we tested the winning model from later work [31] on the data from the same individuals included in the current work. For the RLWM dataset (n=91) [32], we implemented the best known version of the RLWM model [4] with an additional stickiness parameter, which improved model fit significantly. The mathematical formulation of the models can be found in Model equations.

#### 5.2.3. Model-fitting

All models were fitted using the maximum likelihood estimation procedure at the individual level using the MATLAB global optimization toolbox with the fmincon function. Although hierarchical Bayesian methods may have yielded better model fit, we chose to use maximum likelihood estimation because it is simple, efficient, and suffices for our purpose of demonstrating the comparison between the static and dynamic noise models. In practice, we advise users of our dynamic noise estimation method to apply the fitting procedure with the most appropriate assumptions for the model and data.

#### 5.2.4. Model validation and recovery

In model validation, we simulated choice behavior for each subject repeatedly (e.g., for 100 times) using the maximum likelihood parameters obtained from model-fitting. For simulations with dynamic noise estimation, we used the latent state probability – *p*(*Random*) and *p*(*Engaged*) – trajectories inferred from real data to simulate latent state occupancy. To validate how well the models captured behavior, we compared behavioral signatures (e.g., learning curves) between these model simulations and the data (real or simulated) that the models were fitted to.

The recovery of the occupancy probabilities of model latent states was performed by simulating data 30 times per individual using best fit parameters and inferring occupancy probabilities from these data. Model parameters were recovered by first simulating behavior using best fit parameters and refitting the model to the simulated behavior to estimate parameter values. All recovery was performed at the individual level.

## 6. Data and code availability

All data and code used to produce figures in this manuscript can be downloaded at: https://osf.io/b9tmn/?view_only=ba4e06cd8bc8475a8fe131561459f299

## 7. Acknowledgements

This work was supported by the NIH Grant 1R01MH119383.

## Appendix A. Supplementary figures

**Figure A.7:**
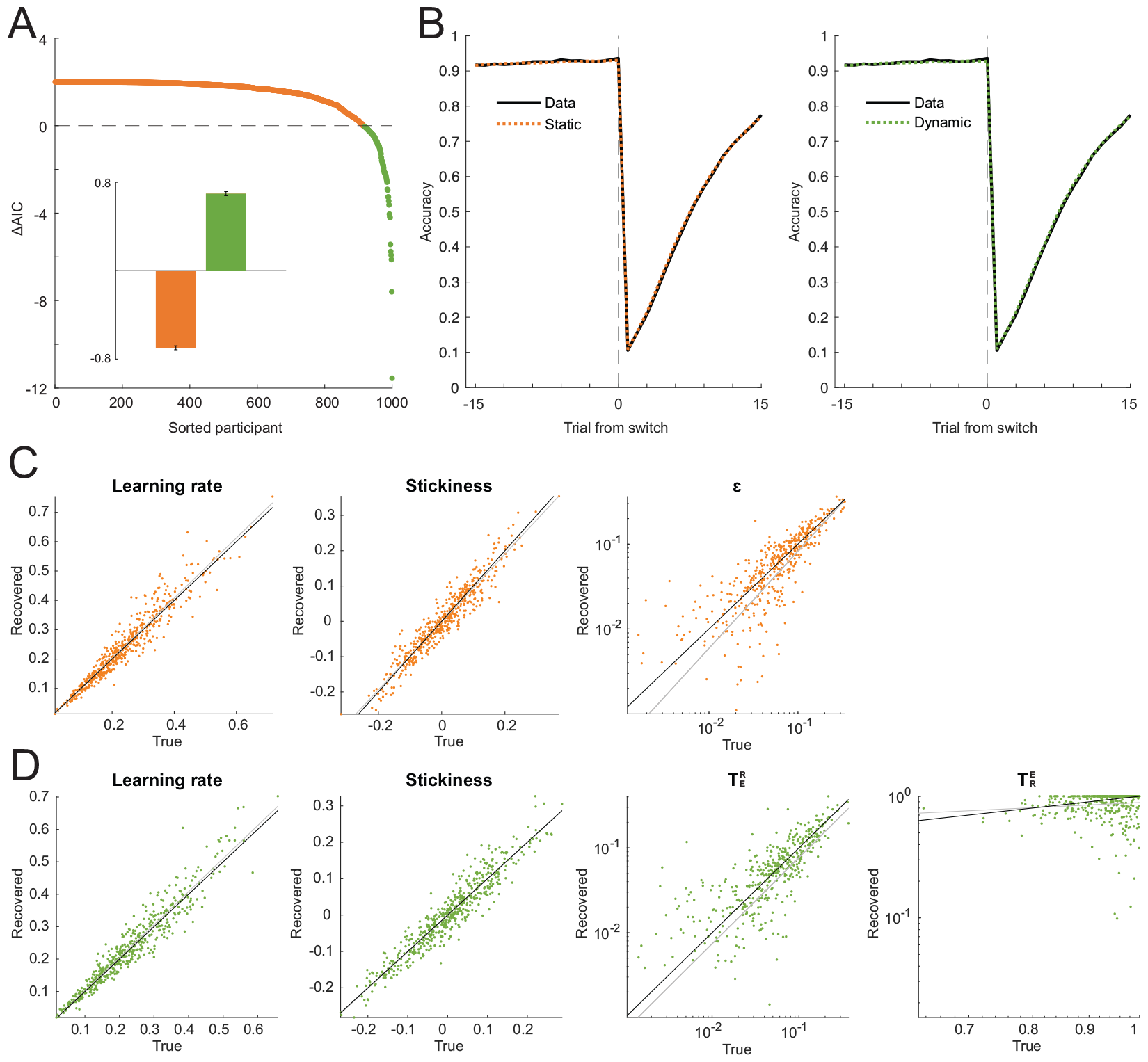
Both models with static and dynamic noise estimation can fully capture behavior and recover generative parameter values when the true model has static noise. A: Evaluation of model fit with AIC on the data of 1,000 participants simulated using the static noise model. Each dot shows the difference in AIC for an individual between the static and dynamic models. A positive value (orange) indicates that the static model is favored and a negative value (green) means that the dynamic model is preferred by the criterion. The inset shows the mean difference in AIC between the models at the group level. B: Learning curves of both models and data. C: Parameter recovery using the static model. D: Parameter recovery using the dynamic model. For the dynamic equivalent of the static model, 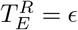 and 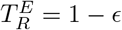.

**Figure A.8:**
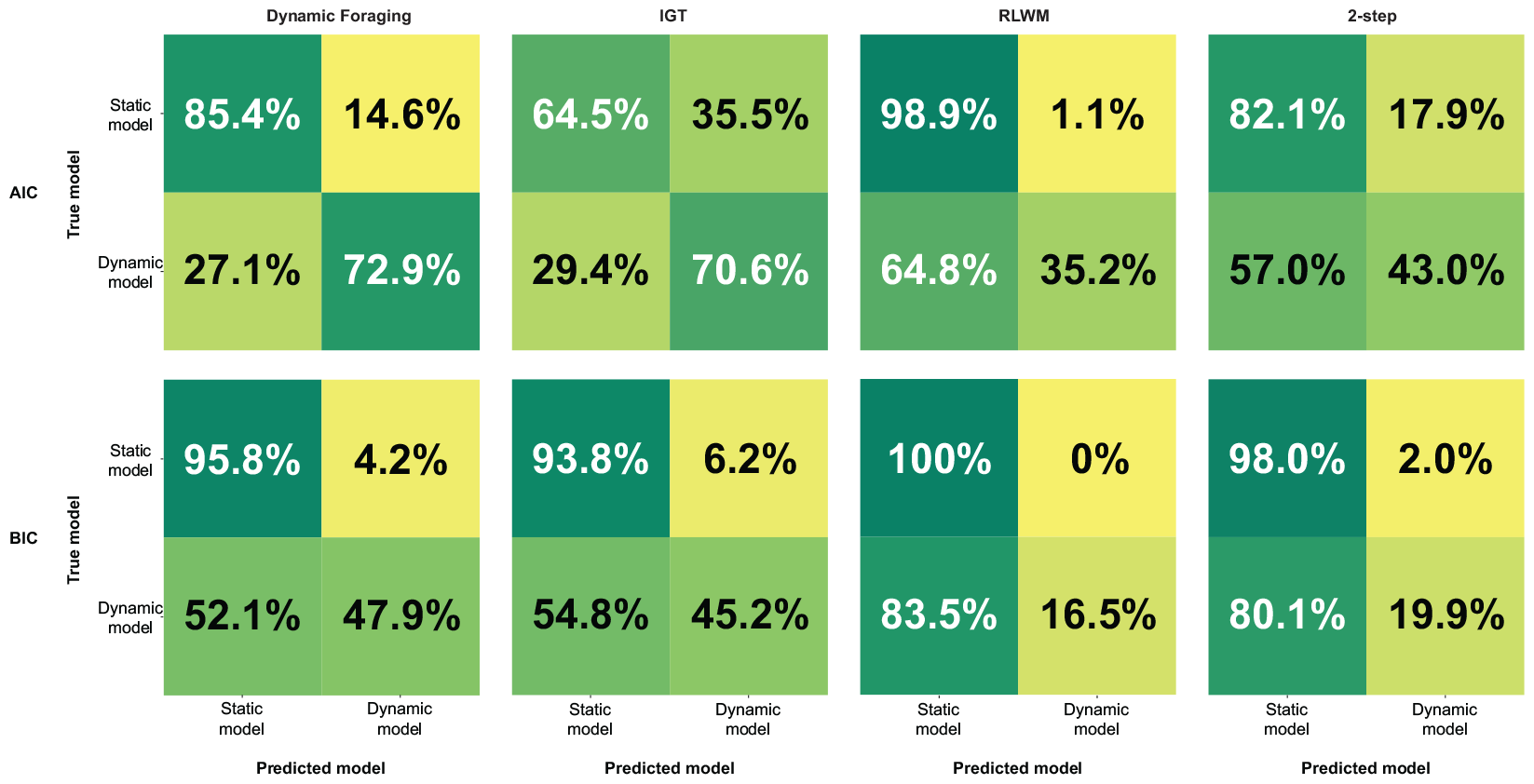
Model identification using AIC and BIC. We performed model identification validation with confusion matrices [2]. To do so, we simulated data with parameters fitted to subjects’ data. The AIC metric yielded better model identification than BIC. We note that simulations of the dynamic noise model were often mis-classified as being generated by the static noise model in RLWM and 2-step datasets. This is because most subjects in these datasets did not benefit substantially from dynamic noise estimation, and the parameters inferred made the dynamic noise model very similar to the static noise model. Thus, simulated behavior was in a range where both models were indistinguishable (since the static noise model is nested in the dynamic one). In these cases, the trivial improvements on likelihoods would be insufficient to offset the penalty incurred by the extra parameter in the dynamic model.

**Figure A.9:**
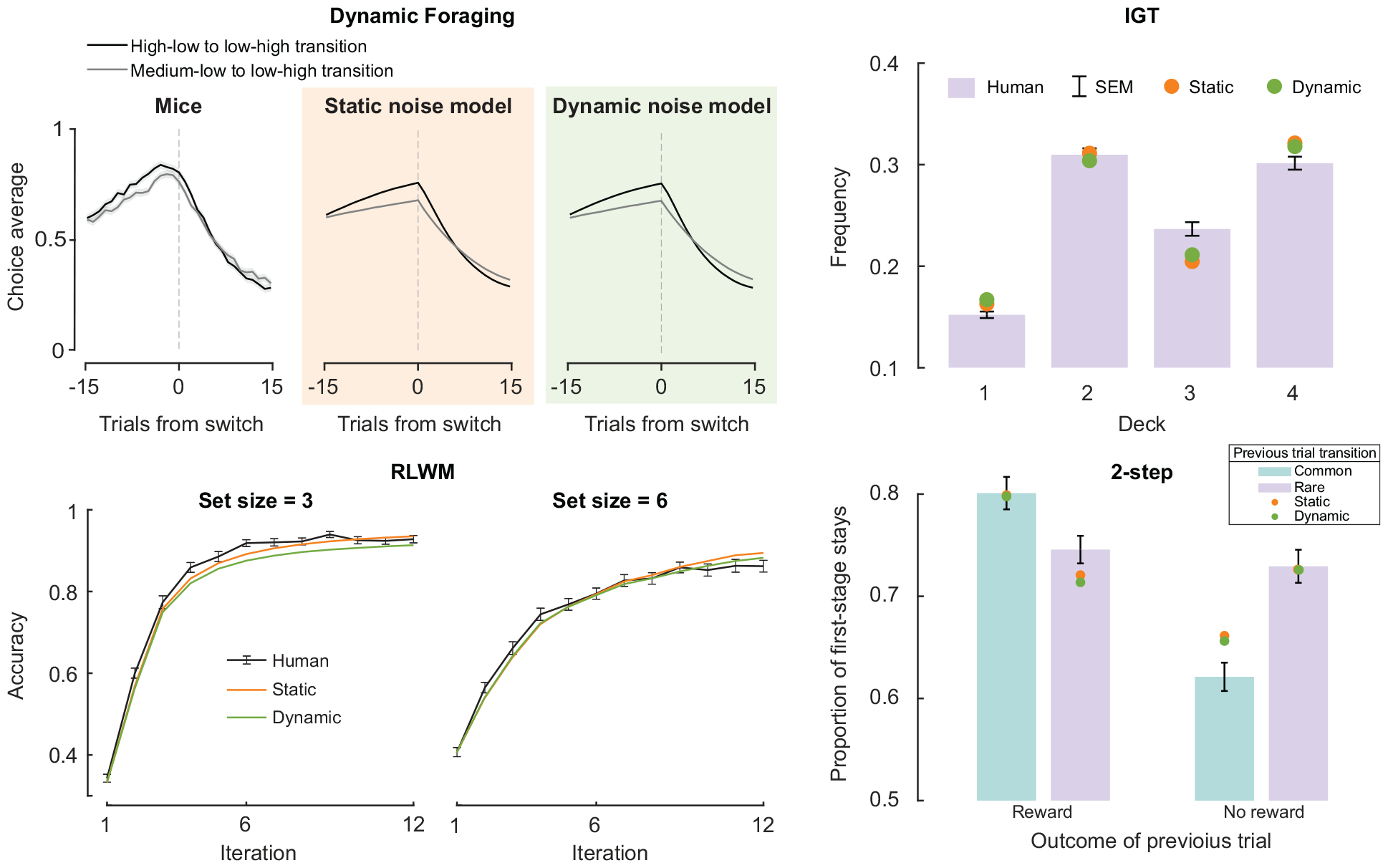
Model validation results on the empirical datasets. Dynamic noise estimation did not alter the qualitative behavioral predictions made by the models.

**Figure A.10:**
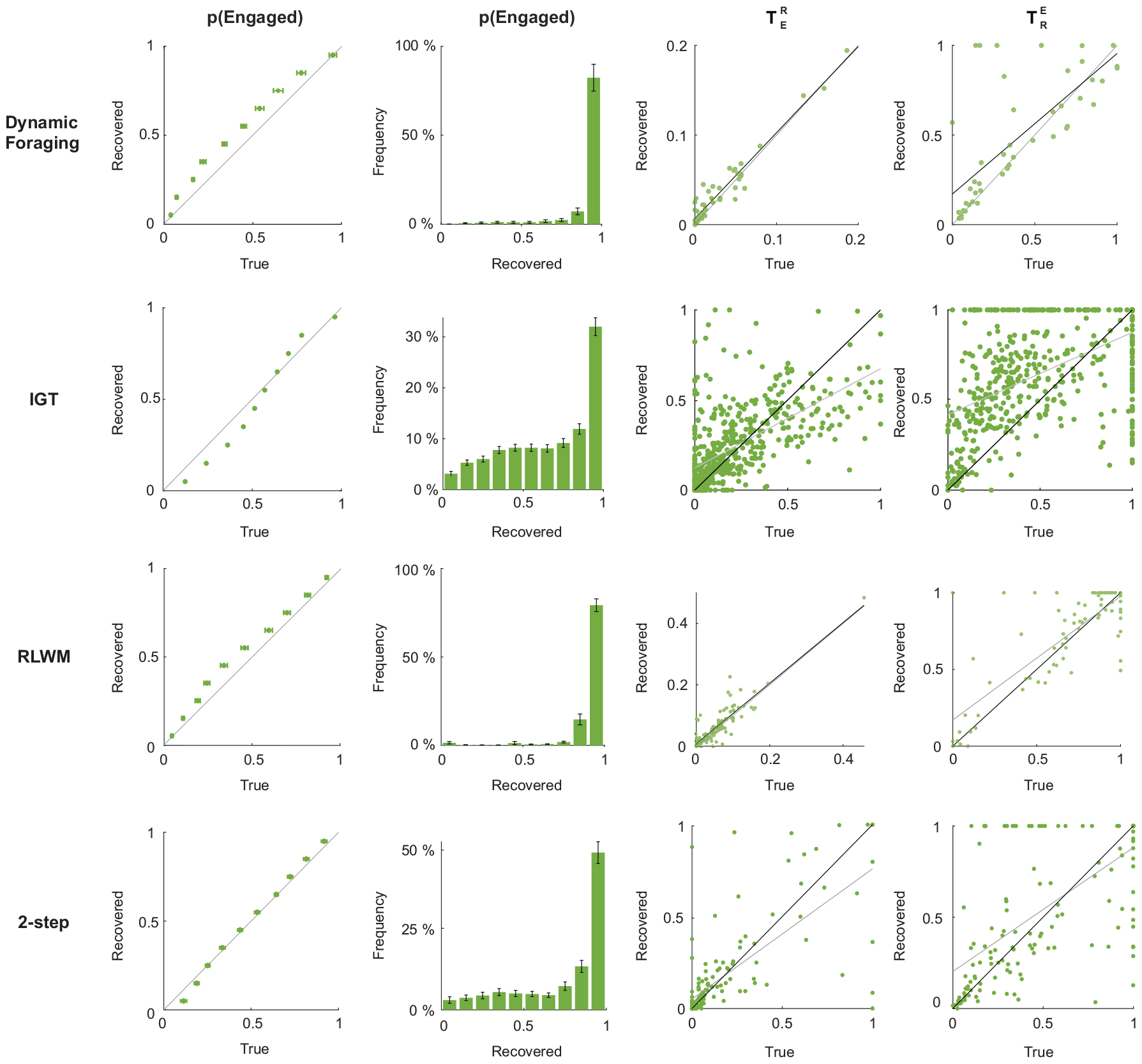
Recovery of latent state occupancy probability and noise parameters. *p*(*Engaged*) recovered well across datasets, with most recovered values between 0.9 and 1. 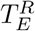 recovery was robust overall, while 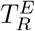 recovered inadequately. This is because the lack of data in the random state led to insufficient potential transitions from the random to engaged state, which under-powered 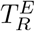 recovery.

**Figure A.11:**
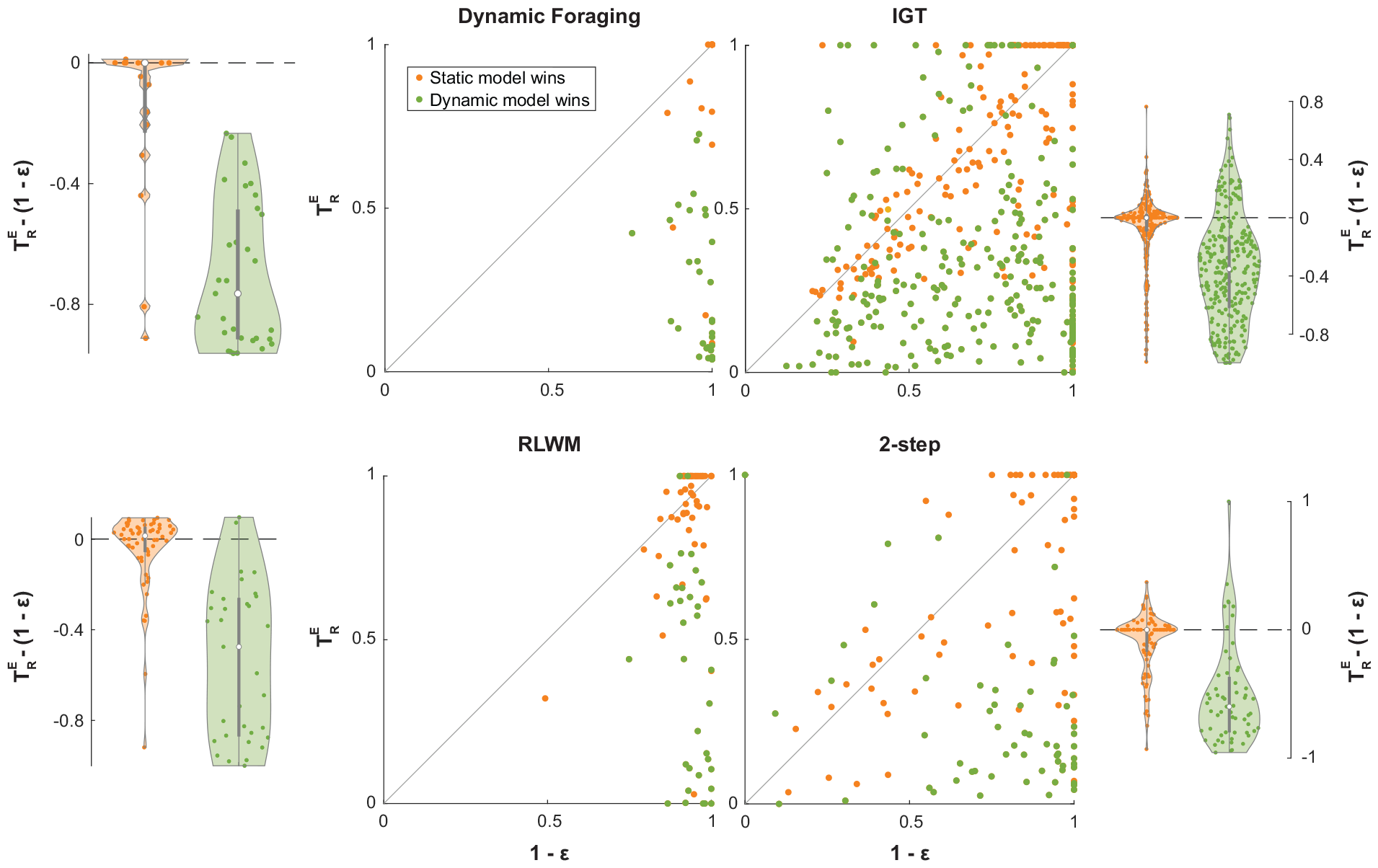
Improved fit by dynamic noise estimation is correlated to decreased estimation of the transition probability from the the random to engaged state.

**Figure A.12:**
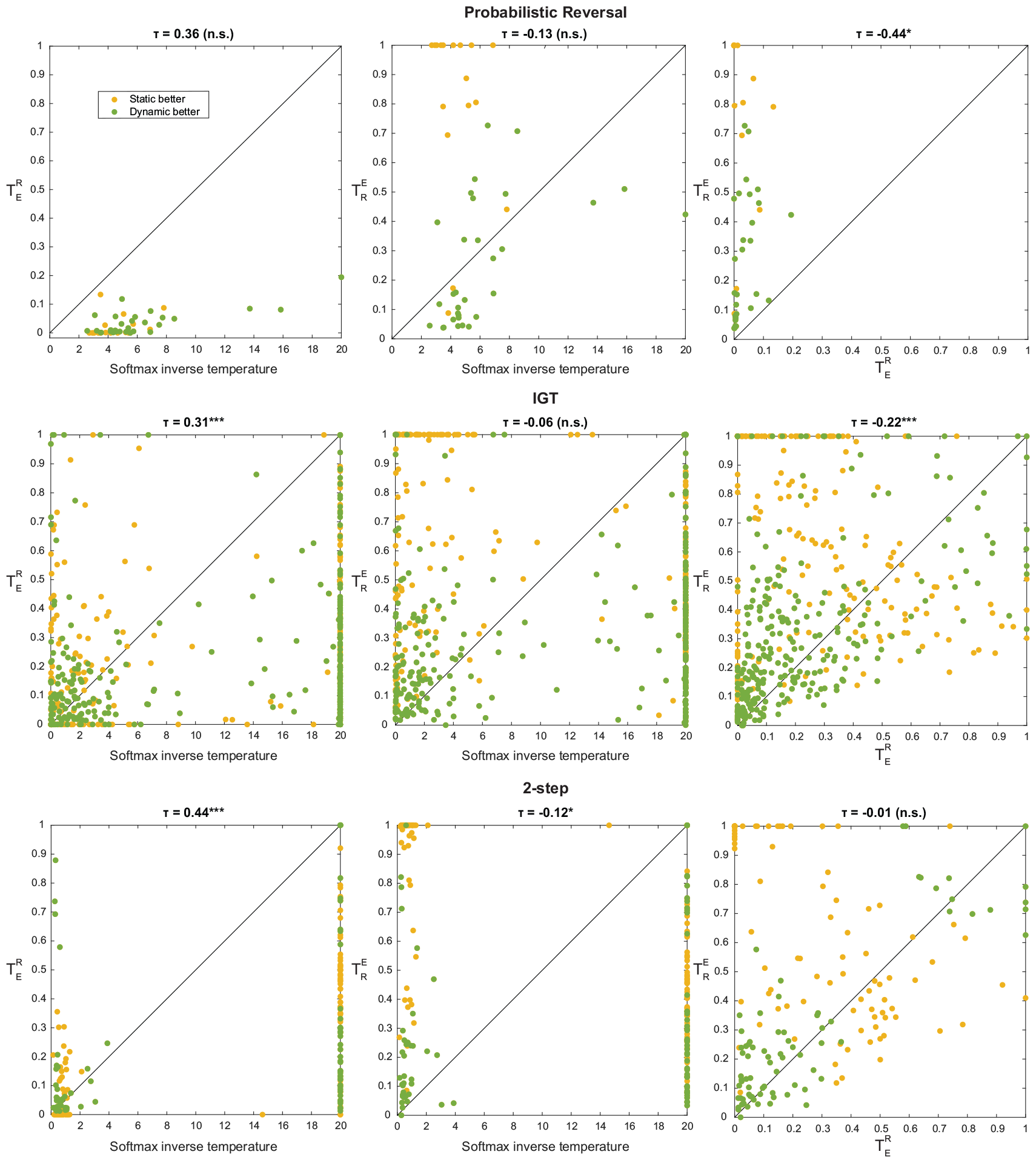
Relationships between noise parameters on the Dynamic Foraging [29], IGT [30], and 2-step [36] datasets. No consistent correlations were found between the noise parameters including the softmax inverse temperature, 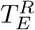, and 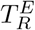.

**Figure A.13:**
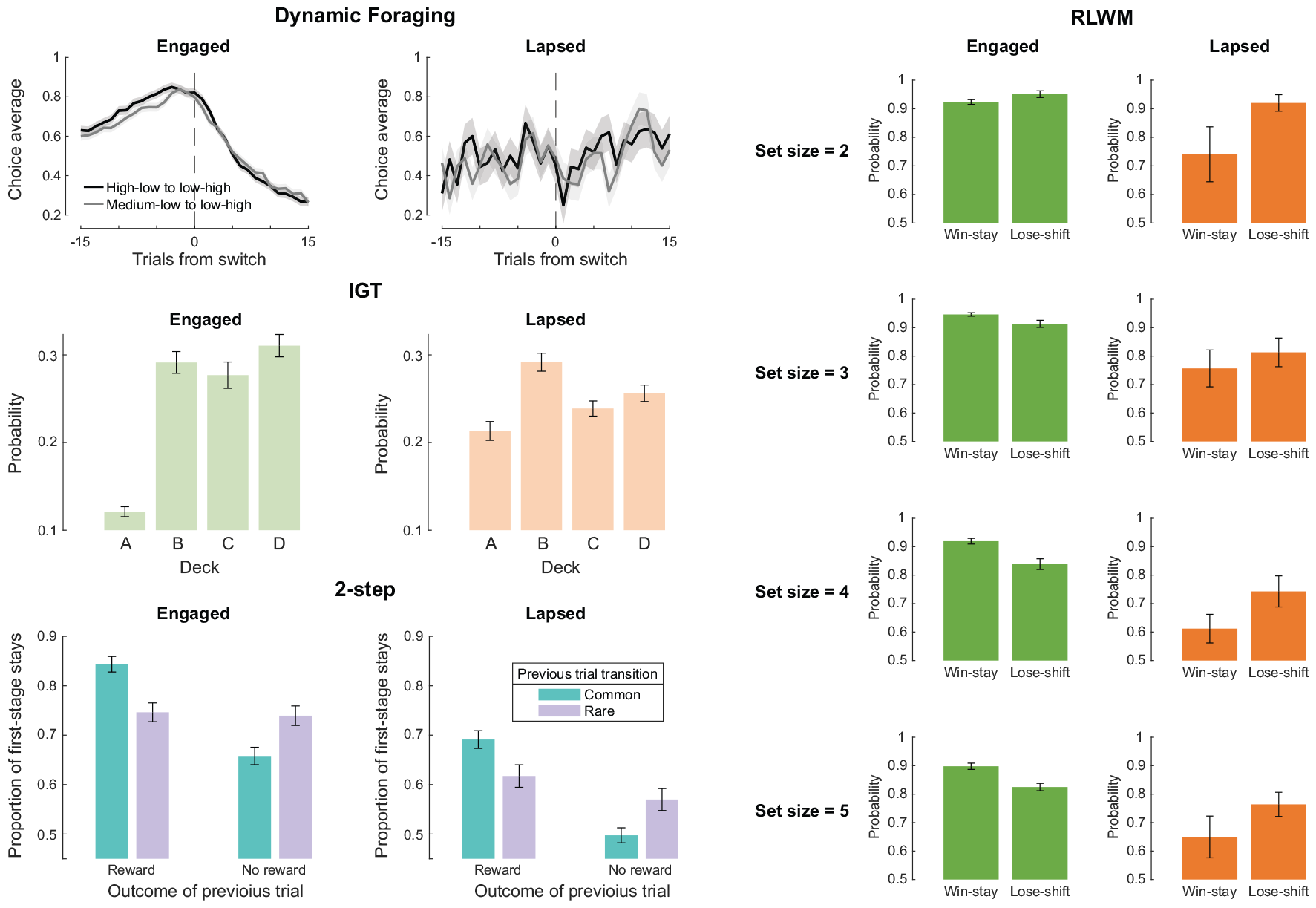
Behavior on putative engaged and lapsed trials predicted by dynamic noise estimation on the Dynamic Foraging [29], IGT [30], 2-step [33], and RLWM [4, 5] datasets. On Dynamic Foraging, the learning curves around switches appear random-like during putative lapses. On the IGT dataset, choice frequencies of decks A and D regressed to the random level (one-tailed Wilcoxon signed-rank test *p* = 9.35*×*10^*-*20^ for A, *p* = 0.48 for B, *p* = 0.11 for C, and *p* = 2.83 *×* 10^*-*5^ for D). For 2-step, the accuracy decreased for all trial types (one-tailed Wilcoxon signed-rank test *p* = 1.73 *×* 10^*-*5^ for common and rewarded previous trials, *p* = 0.019 for rare and rewarded previous trials, *p* = 5.33 *×* 10^*-*4^ for common and unrewarded previous trials, and *p* = 0.002 for rare and unrewarded previous trials). On the RLWM dataset, the win-stay probability decreased more than the lose-shift probability overall (set size of 2: *p* = 0.056 for win-stay and *p* = 0.38 for lose-shift; set size of 3: *p* = 0.07 for win-stay and *p* = 0.092 for lose-shift; set size of 4: *p* = 2.9 *×* 10^*-*4^ for win-stay and *p* = 0.34 for lose-shift; set size of 5: *p* = 0.006 for win-stay and *p* = 0.28 for lose-shift).

**Figure A.14:**
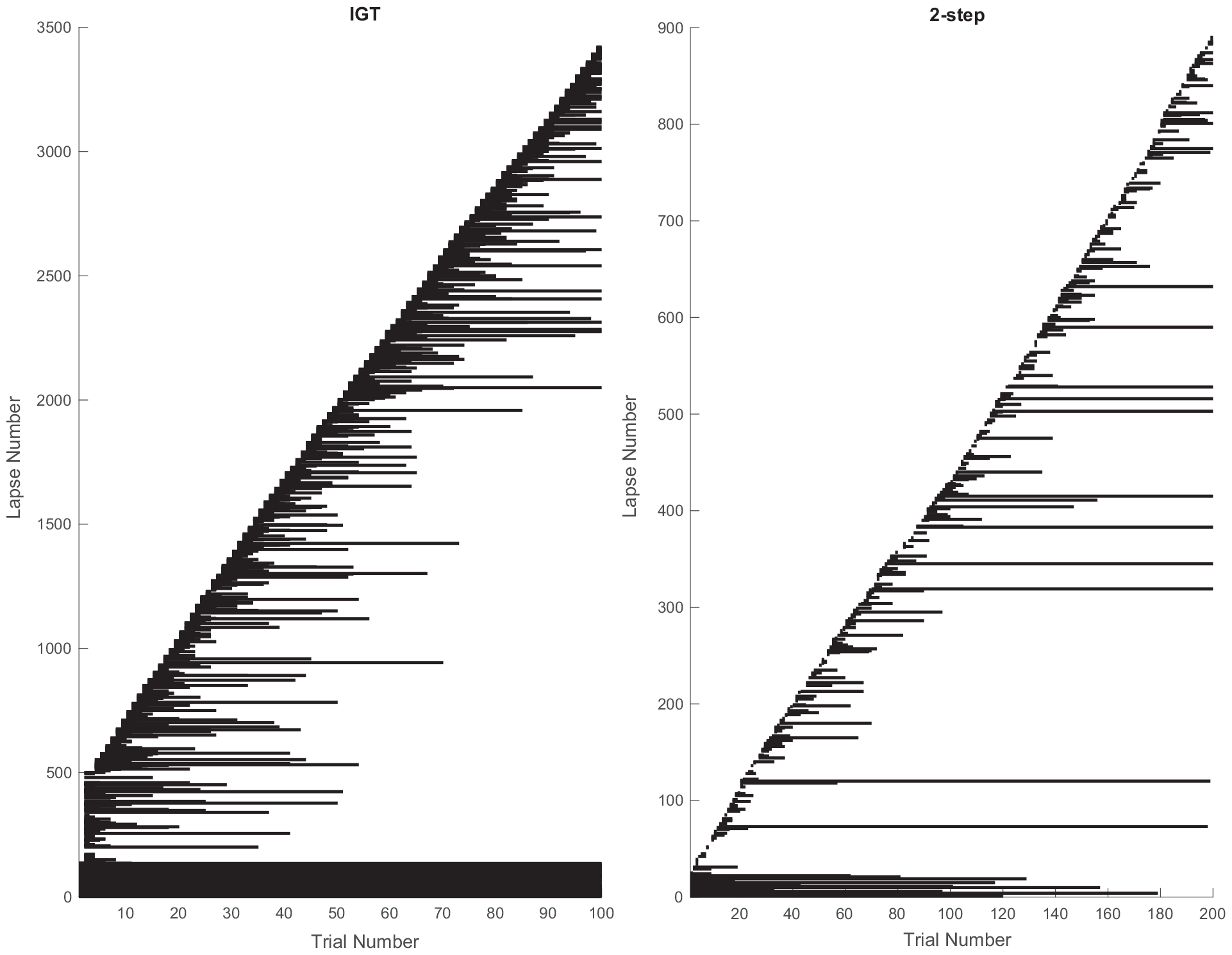
Putative lapses identified by dynamic noise estimation on the IGT [30] and 2-step [33] datasets, both with fixed numbers of trials across participants. The lapses were identified as trials with *p*(*Engaged*) *<* 0.5, sorted by the start trial, and shown across participants.

**Figure A.15:**
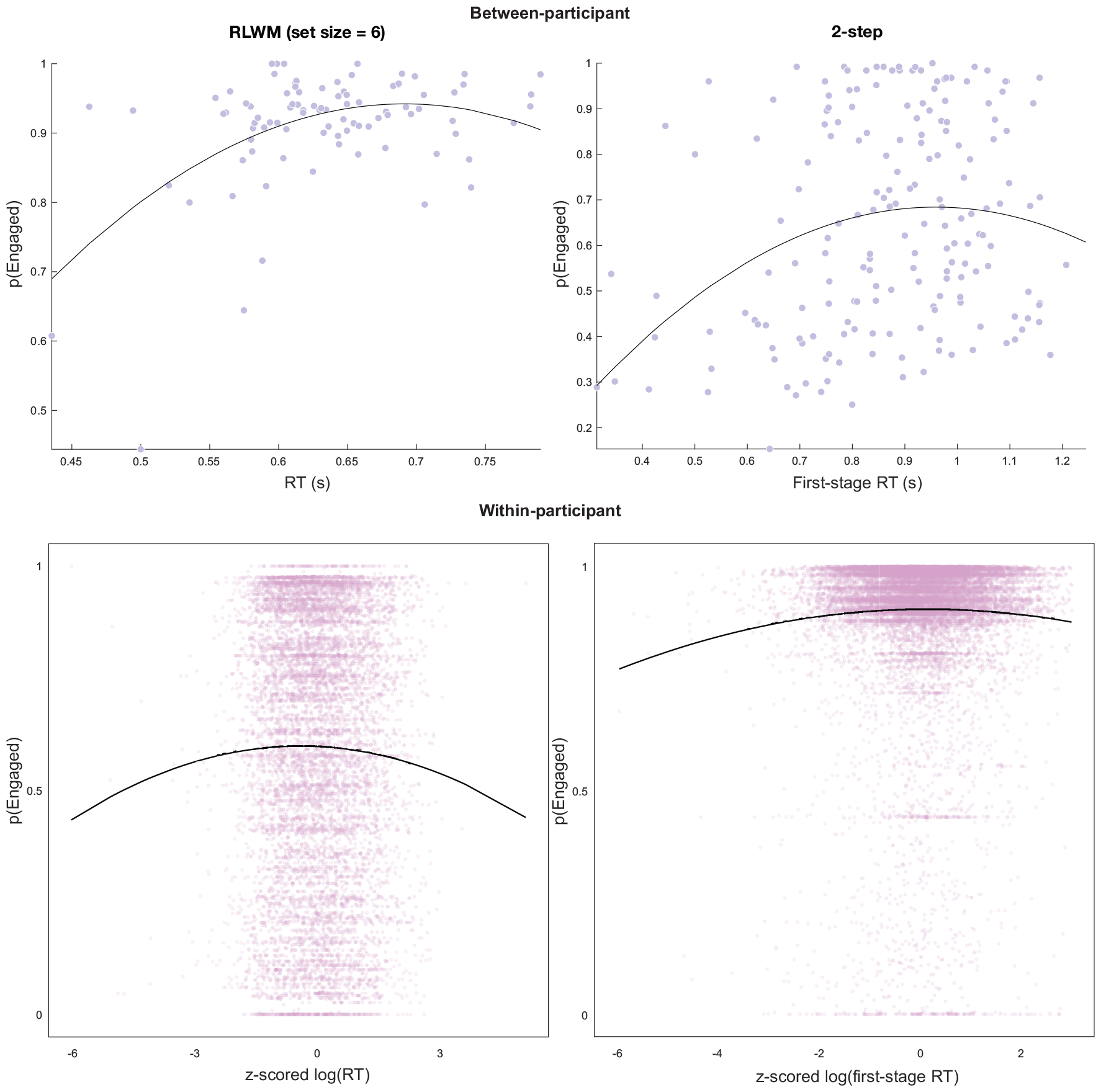
The inverted-U relationship between *p*(*Engaged*) and reaction time between- and within-participants on the RLWM [32] and 2-step [36] datasets. All p-values are less than 0.01 for the regression coefficients of the quadratic terms. The specific statistics are reported in Results.

**Figure A.16:**
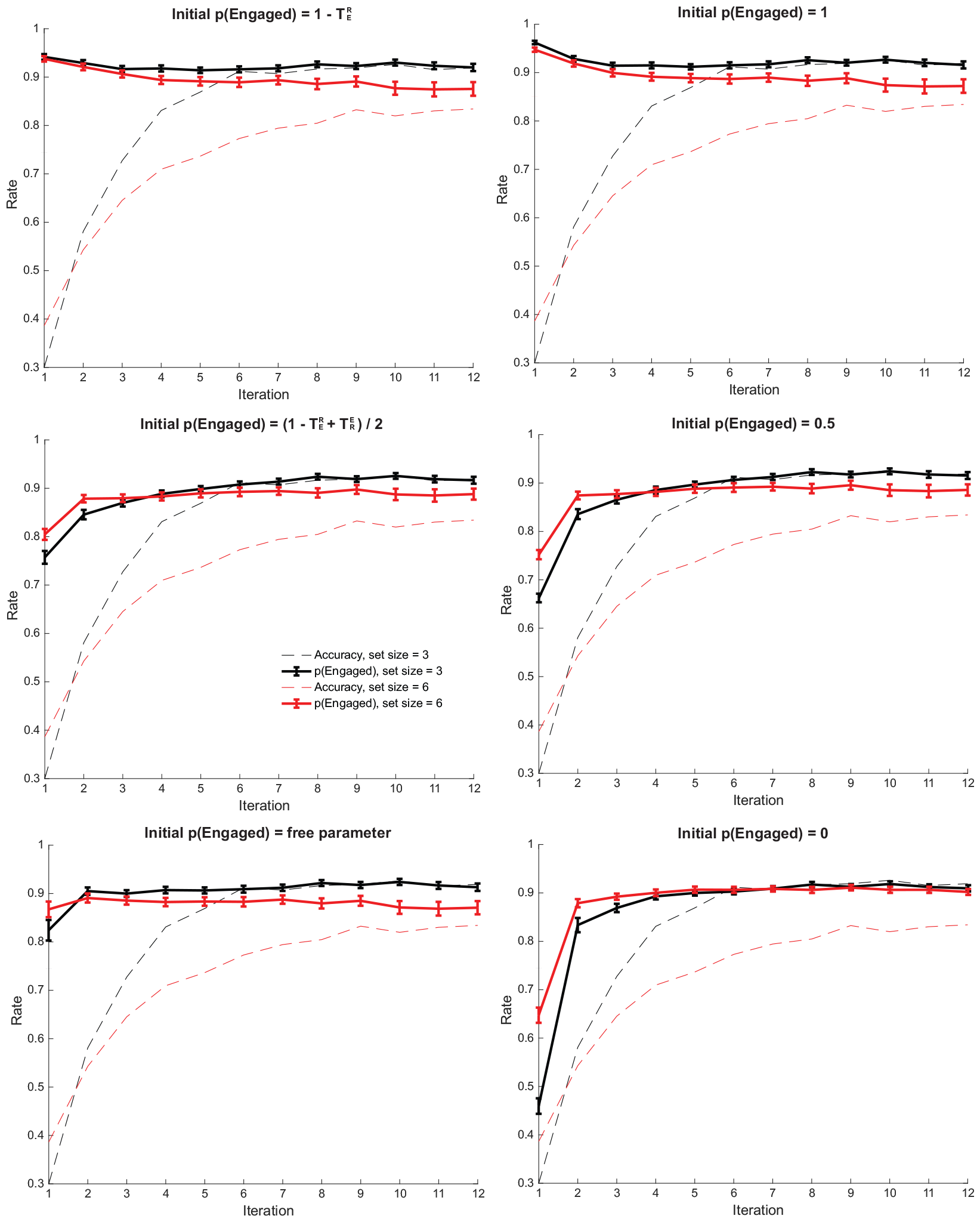
Different ways to initialize *p*(*Engaged*) lead to different latent state occupancy estimations in the first few trials, but similar trajectories afterwards. Note that the estimated engaged probability does not always follow the same trend as accuracy: towards the end of the block, while the difference in accuracy between set sizes of 3 and 6 shrinks, the difference in *p*(*Engaged*) does not.

## Appendix B. Model equations

### Appendix B.1. Probabilistic Reversal

The model for the Probabilistic Reversal environment consists of 2 free parameters: *α* (learning rate) and *ϕ* (choice stickiness). The softmax inverse temperature is fixed at, *β* = 8.

On trial *t*, the choice is made according to action probabilities computed through the softmax function. For example, the probability of choosing the left action is:

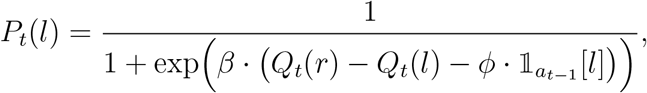

where 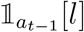 takes on the value of 1 if *a*_*t-*1_ = *l* and -1 otherwise.

Once the reward r_*t*_ has been observed, the action values are updated:

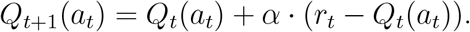

### Appendix B.2. Dynamic Foraging

The meta-learning model in the original paper was implemented [29]. The model has 7 parameters:, *β* (softmax inverse temperature), bias (for the right action), *α*_(+)_ (positive learning rate), 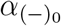 (baseline negative learning rate), *α*_*υ*_ (rate of RPE magnitude integration), *ξ* (meta-learning rate for unexpected uncertainty), and ∼ (forgetting rate).

On trial *t*, a decision is sampled from choice probabilities obtained through a softmax decision function applied to the action values of the left and right actions:

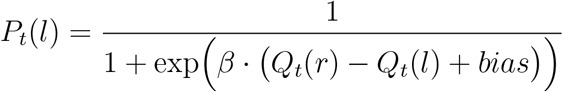

and

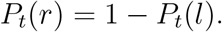

Once the reward is observed, assuming the left action is chosen, its value is updated as follows:

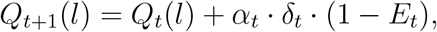

where *α*_*t*_ is *α*_(+)_ if the reward-prediction error (RPE), *δ* _*t*_ = *R*_*t*_ *-Q*_*t*_(*l*), is positive, and 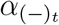 otherwise. *E*_*t*_ is an evolving estimate of expected uncertainty calculated from the history of absolute RPEs:

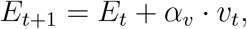

where

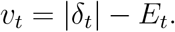

When the RPE is negative, the negative learning rate is dynamically adjusted and lower-bounded by 0:

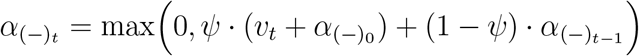

Finally, the unchosen action (e.g., right) is forgotten:

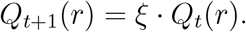

### Appendix B.3. IGT

The Value plus Sequential Exploration model [31] was implemented for the IGT dataset. The model is defined by 5 parameters: *α* (learning rate), *β* (softmax inverse temperature), *θ* (value sensitivity), Δ (decay), and *ϕ* (exploration bonus).

On trial *t*, the decision is sampled based on the probability of choosing deck *d*:

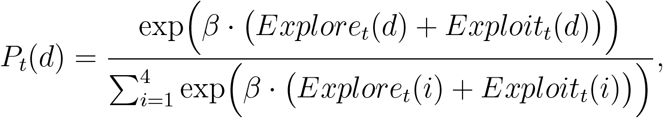

where Explore_*t*_(*d*) and Exploit_*t*_(*d*) are the action values of deck d using the exploration and exploitation weights. For the selected deck, their values are updated according to the following equations:

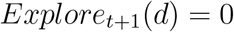

and

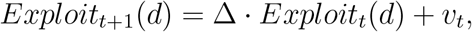

where υ _*t*_ = (*Gain*_*t*_)^*θ*^ *-* (*Loss*_*t*_)^*θ*^. For the unselected decks, the weights are controlled by the following equations:

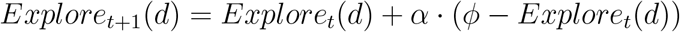

and

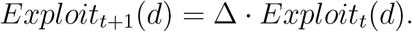

### Appendix B.4. RLWM

The RLWM model is improved upon previously published versions [4, 32] by the inclusion of a choice stickiness parameter. The model has 6 parameters in total: *α* (learning rate), *bias* (for negative learning), *ϕ* (stickiness), *ρ* (working memory weight), *γ* (forgetting rate), and *K* (working memory capacity). The softmax inverse temperature parameter is fixed at, *β* = 20.

On trial *t*, the probability of choosing an action *a*_*t*_ in state *s*_*t*_ is given by a weighted combination between a reinforcement learning policy and a working memory one:

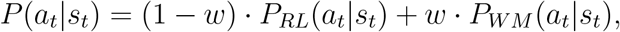

where 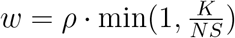 and *NS* is the set size. The action values for both policies are computed as follows:

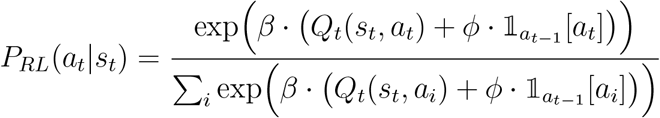

and

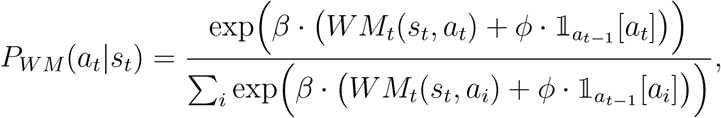

where 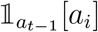 is an indicator that takes on the value of 1 if 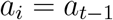 and 0 otherwise.

All working memory values are forgotten on each trial:

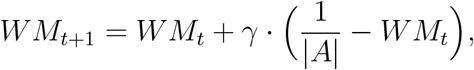

where |*A*| is the total number of available actions. The values are then updated according to the following equations:

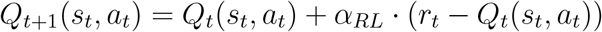

and

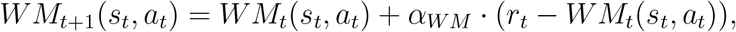

where if *r*_*t*_ = 1, *α*_*RL*_ = *α* and *α*_*WM*_ = 1, and if *r*_*t*_ = 0, *α*_*RL*_ = *bias · α* and α_*WM*_ = *bias*.

### Appendix B.5. 2-step

The 2-step model [33] contains 6 free parameters: *α* (learning rate), *β*_*MB*_ (softmax inverse temperature for the model-based policy), *β*_*MF*_ (softmax inverse temperature for the model-free policy), *β*(softmax inverse temperature for the second stage), *p* (stimulus stickiness), and *ϕ* (response stickiness).

The first-stage decision is made according to action probabilities computed using both the model-based and model-free action values:

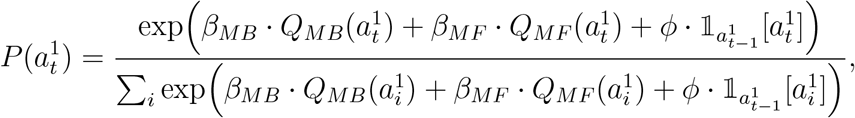

where 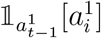 is an indicator that takes on the value of 1 if 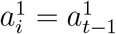 and 0 otherwise. The second-stage action probabilities are also computed through the softmax function:

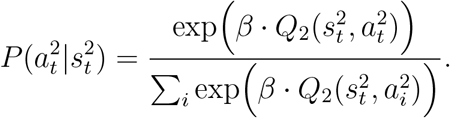

Once the reward *r*_*t*_ has been observed, the action values are updated as follows:

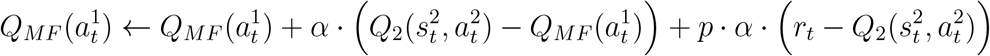

and

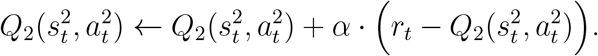

Note that the model-based action values do not need to be updated and can be computed directly:

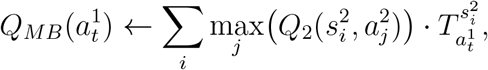

where 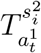 is the transition probability from the first-stage choice 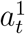 to the second-stage state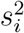, which the agent is assumed to know.

## References

[1] Palminteri, S., Wyart, V. & Koechlin, E. The importance of falsification in computational cognitive modeling. Trends In Cognitive Sciences. 21, 425–433 (2017)

[2] Wilson, R. & Collins, A. Ten simple rules for the computational modeling of behavioral data. Elife. 8 pp. e49547 (2019)

[3] Kass, R. & Raftery, A. Bayes factors. Journal Of The American Statistical Association. 90, 773–795 (1995)

[4] Master, S., Eckstein, M., Gotlieb, N., Dahl, R., Wilbrecht, L. & Collins, A. Disentangling the systems contributing to changes in learning during adolescence. Developmental Cognitive Neuroscience. 41 pp. 100732 (2020)

[5] Eckstein, M., Master, S., Xia, L., Dahl, R., Wilbrecht, L. & Collins, A. The interpretation of computational model parameters depends on the context. Elife. 11 pp. e75474 (2022)

[6] Lee, M. & Webb, M. Modeling individual differences in cognition. Psychonomic Bulletin & Review. 12, 605–621 (2005)

[7] Huys, Q., Browning, M., Paulus, M. & Frank, M. Advances in the computational understanding of mental illness. Neuropsychopharmacology. 46, 3–19 (2021)

[8] Tversky, A. & Kahneman, D. Advances in prospect theory: Cumulative representation of uncertainty. Journal Of Risk And Uncertainty. 5, 297–323 (1992)

[9] Bitzer, S., Park, H., Blankenburg, F. & Kiebel, S. Perceptual decision making: drift-diffusion model is equivalent to a Bayesian model. Frontiers In Human Neuroscience. 8 pp. 102 (2014)

[10] Dayan, P. & Niv, Y. Reinforcement learning: the good, the bad and the ugly. Current Opinion In Neurobiology. 18, 185–196 (2008)

[11] Esterman, M. & Rothlein, D. Models of sustained attention. Current Opinion In Psychology. 29 pp. 174–180 (2019)

[12] Warm, J., Parasuraman, R. & Matthews, G. Vigilance requires hard mental work and is stressful. Human Factors. 50, 433–441 (2008)

[13] Wilson, R., Geana, A., White, J., Ludvig, E. & Cohen, J. Humans use directed and random exploration to solve the explore–exploit dilemma. Journal Of Experimental Psychology: General. 143, 2074 (2014)

[14] Findling, C. & Wyart, V. Computation noise in human learning and decision-making: origin, impact, function. Current Opinion In Behavioral Sciences. 38 pp. 124–132 (2021)

[15] Sutton, R. & Barto, A. Reinforcement learning: An introduction. (MIT press,2018)

[16] Chapelle, O. & Li, L. An empirical evaluation of thompson sampling. Advances In Neural Information Processing Systems. 24 (2011)

[17] Thompson, W. On the likelihood that one unknown probability exceeds another in view of the evidence of two samples. Biometrika. 25, 285–294 (1933)

[18] Wang, S. & Wilson, R. Any way the brain blows? The nature of decision noise in random exploration. (PsyArXiv,2018)

[19] Daw, N. & Tobler, P. Value learning through reinforcement: the basics of dopamine and reinforcement learning. Neuroeconomics. pp. 283–298 (2014)

[20] Collins, A. & Frank, M. How much of reinforcement learning is working memory, not reinforcement learning? A behavioral, computational, and neurogenetic analysis. European Journal Of Neuroscience. 35, 1024–1035 (2012)

[21] Nassar, M. & Frank, M. Taming the beast: extracting generalizable knowledge from computational models of cognition. Current Opinion In Behavioral Sciences. 11 pp. 49–54 (2016)

[22] Schaaf, J., Jepma, M., Visser, I. & Huizenga, H. A hierarchical Bayesian approach to assess learning and guessing strategies in reinforcement learning. Journal Of Mathematical Psychology. 93 pp. 102276 (2019)

[23] Yechiam, E. & Busemeyer, J. Comparison of basic assumptions embedded in learning models for experience-based decision making. Psychonomic Bulletin & Review. 12, 387–402 (2005)

[24] Schulz, E. & Gershman, S. The algorithmic architecture of exploration in the human brain. Current Opinion In Neurobiology. 55 pp. 7–14 (2019)

[25] Group, T., Fawcett, T., Fallenstein, B., Higginson, A., Houston, A., Mallpress, D., Trimmer, P. & McNamara, J. The evolution of decision rules in complex environments. Trends In Cognitive Sciences. 18, 153–161 (2014)

[26] Fisher, R. On the mathematical foundations of theoretical statistics. Philosophical Transactions Of The Royal Society Of London. Series A, Containing Papers Of A Mathematical Or Physical Character. 222, 309–368 (1922)

[27] Piray, P., Dezfouli, A., Heskes, T., Frank, M. & Daw, N. Hierarchical Bayesian inference for concurrent model fitting and comparison for group studies. PLoS Computational Biology. 15, e1007043 (2019)

[28] Izquierdo, A., Brigman, J., Radke, A., Rudebeck, P. & Holmes, A. The neural basis of reversal learning: an updated perspective. Neuroscience. 345 pp. 12–26 (2017)

[29] Grossman, C., Bari, B. & Cohen, J. Serotonin neurons modulate learning rate through uncertainty. Current Biology. 32, 586–599 (2022)

[30] Steingroever, H., Fridberg, D., Horstmann, A., Kjome, K., Kumari, V., Lane, S., Maia, T., McClelland, J., Pachur, T., Premkumar, P. & Others Data from 617 healthy participants performing the Iowa gambling task: A” many labs” collaboration. Journal Of Open Psychology Data. 3, 340–353 (2015)

[31] Ligneul, R. Sequential exploration in the Iowa gambling task: validation of a new computational model in a large dataset of young and old healthy participants. PLoS Computational Biology. 15, e1006989 (2019)

[32] Collins, A. The tortoise and the hare: Interactions between reinforcement learning and working memory. Journal Of Cognitive Neuroscience. 30, 1422–1432 (2018)

[33] Nussenbaum, K., Scheuplein, M., Phaneuf, C., Evans, M. & Hartley, C. Moving developmental research online: comparing in-lab and web-based studies of model-based reinforcement learning. Collabra: Psychology. 6 (2020)

[34] Akaike, H. A new look at the statistical model identification. IEEE Transactions On Automatic Control. 19, 716–723 (1974)

[35] Rigoux, L., Stephan, K., Friston, K. & Daunizeau, J. Bayesian model selection for group studies—revisited. Neuroimage. 84 pp. 971–985 (2014)

[36] Kool, W., Cushman, F. & Gershman, S. When does model-based control pay off?. PLoS Computational Biology. 12, e1005090 (2016)

[37] Luce, R. Individual choice behavior: A theoretical analysis. (Courier Corporation,2012)

[38] Daw, N., O’doherty, J., Dayan, P., Seymour, B. & Dolan, R. Cortical substrates for exploratory decisions in humans. Nature. 441, 876–879 (2006)

[39] Wilson, R., Geana, A., White, J., Ludvig, E. & Cohen, J. Humans use directed and random exploration to solve the explore–exploit dilemma. Journal Of Experimental Psychology: General. 143, 2074 (2014)

[40] Collins, A. & Koechlin, E. Reasoning, learning, and creativity: frontal lobe function and human decision-making. PLoS Biology. 10, e1001293 (2012)

[41] Donoso, M., Collins, A. & Koechlin, E. Foundations of human reasoning in the prefrontal cortex. Science. 344, 1481–1486 (2014)

[42] Aston-Jones, G., Rajkowski, J. & Cohen, J. Role of locus coeruleus in attention and behavioral flexibility. Biological Psychiatry. 46, 1309–1320 (1999)

[43] Berridge, C. & Waterhouse, B. The locus coeruleus–noradrenergic system: modulation of behavioral state and state-dependent cognitive processes. Brain Research Reviews. 42, 33–84 (2003)

[44] Ashwood, Z., Roy, N., Stone, I., Laboratory, I., Urai, A., Churchland, A., Pouget, A. & Pillow, J. Mice alternate between discrete strategies during perceptual decision-making. Nature Neuroscience. 25, 201–212 (2022)

[45] Trach, J., DeBettencourt, M., Radulescu, A. & McDougle, S. Reward prediction errors modulate attentional vigilance. (PsyArXiv,2022)

[46] Botvinick, M., Braver, T., Barch, D., Carter, C. & Cohen, J. Conflict monitoring and cognitive control. Psychological Review. 108, 624 (2001)

[47] Laeng, B., Sirois, S. & Gredebäck, G. Pupillometry: A window to the preconscious?. Perspectives On Psychological Science. 7, 18–27 (2012)

[48] Polich, J. Updating P300: an integrative theory of P3a and P3b. Clinical Neurophysiology. 118, 2128–2148 (2007)

[49] Huys, Q., Maia, T. & Frank, M. Computational psychiatry as a bridge from neuroscience to clinical applications. Nature Neuroscience. 19, 404–413 (2016)

[50] Barkley, R. Behavioral inhibition, sustained attention, and executive functions: constructing a unifying theory of ADHD. Psychological Bulletin. 121, 65 (1997)

[51] Auer, P., Cesa-Bianchi, N. & Fischer, P. Finite-time analysis of the multiarmed bandit problem. Machine Learning. 47 pp. 235–256 (2002)

[52] Puterman, M. L. (2014). Markov decision processes: discrete stochastic dynamic programming. John Wiley & Sons.

